# Tau regulates Arc stability in neuronal dendrites via a proteasome-sensitive but ubiquitin-independent pathway

**DOI:** 10.1101/2022.12.15.520620

**Authors:** Dina W. Yakout, Ankit Shroff, Vishrut Thaker, Zachary D. Allen, Taras Y. Nazarko, Angela M. Mabb

**Affiliations:** Neuroscience Institute, Georgia State University, Atlanta, Georgia, United States.; Department of Biology, Georgia State University, Atlanta, Georgia, United States.; Center for Behavioral Neuroscience, Atlanta, Georgia, United States.

**Author notes:** Correspondence: Dr. Angela M. Mabb Georgia State University 100 Piedmont Ave. SE Atlanta, GA 30303 USA.

**Keywords:** Arc, tau, AMPA receptors, proteasome, ubiquitination, Alzheimer’s disease

## Abstract

Tauopathies are neurodegenerative disorders characterized by the deposition of aggregates of the microtubule associated protein tau, a main component of neurofibrillary tangles. Alzheimer’s disease (AD) is the most common type of tauopathy and dementia, with amyloid-beta pathology as an additional hallmark feature of the disease. Besides the role of tau in stabilizing microtubules, it is localized at postsynaptic sites and can disrupt synaptic plasticity when knocked out or overexpressed. The activity-regulated cytoskeleton-associated protein (Arc), is an immediate early gene that plays a key role in synaptic plasticity, learning and memory. Arc has been implicated in AD pathogenesis, where it was found to regulate activity-dependent release of amyloid-beta (Aβ). Here we show that Arc protein is upregulated in the hippocampus of tau knockout (Tau KO) mice and in dendrites of Tau KO primary hippocampal neurons. Conversely, overexpression of tau decreased Arc stability exclusively in neuronal dendrites and was coupled to an increase in the expression of dendritic and somatic surface GluA1-containing α-amino-3-hydroxy-5-methyl-4-isoxazolepropionic acid (AMPA) receptors. The Tau-dependent decrease in Arc was proteasome sensitive, yet independent of Arc ubiquitination and required the endophilin-binding domain of Arc, which is essential for promoting the endocytosis of AMPA receptors. Importantly, these effects on Arc stability and GluA1 localization were not observed in the commonly studied tau mutant, P301L. Our findings show a physiological role for tau in regulating Arc and implicate specific variants of tau in regulating Arc stability and AMPA receptor targeting, which may in part explain observed deficits in synaptic plasticity in select types of tauopathies.

## Introduction

Tauopathies are a diverse group of neurodegenerative disorders predominantly characterized by dementia or degeneration of the motor system (1). A hallmark of tauopathies is the accumulation of tau into insoluble aggregates and filaments which is a major component of neurofibrillary tangles (NFTs) in the brain (2,3). Besides tau pathology, tauopathies may also involve other pathological changes such as amyloid deposition that is observed in Alzheimer’s disease (AD) and Down’s syndrome (1). In AD, the most common tauopathy, the progress of tau pathology, follows a stereotypical pattern in the brain that is highly correlated with the progress of cognitive impairment, which led Braak and Braak to base the staging of AD on the pattern of NFT deposition in the brain (4).

Tau is encoded by the *MAPT* gene on chromosome 17 (5). Its C-terminal region contains 18-residue repeats, which together form the microtubule binding domain (MTBD), which is linked to the N-terminal region through a proline-rich region (6). The *MAPT* gene consists of 16 exons, 11 of which are expressed in the central nervous system (7). In humans, six different isoforms of tau have been reported with differences in alternative mRNA splicing of exons *2*, *3* and *10*. Alternative splicing of exons *2* and *3* yields 0,1 or 2 N-terminal repeats (0N, 1N and 2N isoforms) while alternative splicing of exon *10* leads to the presence or absence of the R2 domain, which is one of the 4 repeats that bind to microtubules (3R and 4R isoforms) (8). Tau also undergoes several posttranslational modifications including, but not limited to, phosphorylation (9), acetylation (10,11) and ubiquitination (12). During early stages of development, tau is highly phosphorylated compared to the adult brain (13). In tauopathies, tau becomes hyperphosphorylated, which is thought to increase its propensity to form aggregates and reduce its affinity for microtubules (14). Additionally, it has been shown that some degree of tau accumulation and hyperphosphorylation occurs in normal aging (15,16).

There are over 50 mutants of the *MAPT* gene that have been identified in several tauopathies (17). Some of these mutations can affect the alternative splicing of *tau* mRNA leading to overproduction of 3R or 4R isoforms and thus pathologically increasing tau and facilitating its aggregation (18). Other mutations, like the missense P301L mutation found within the 2R region, increase tau phosphorylation and decrease its binding to microtubules resulting in increased levels of free tau, which is thought to promote its aggregation (19).

With tau as a key molecular player in AD and tauopathies, understanding the physiological role of tau is crucial for understanding its role in pathological conditions and the downstream effects of the loss or gain of tau function. Over the past decade, multiple studies have focused on physiological and pathological roles for tau beyond those related to microtubule stabilization (21). Tau is enriched in neuronal axons, with lower levels detected in the plasma membrane, dendrites and dendritic spines, with a differential spatial distribution of tau isoforms (22). Studies from Tau KO mice have shown that loss of tau does not lead to gross behavioral or neuronal changes in young mice. However; tau does modulate synaptic plasticity, where Tau KO mice have deficits in long-term potentiation (LTP) and long-term depression (LTD) (23,24).

Characterization of the tau interactome in the mouse brain identified proteins involved in synaptic vesicle cycling and postsynaptic receptor trafficking (25). Yet, a mechanism for the physiological role of tau in regulating synaptic plasticity has not been clearly elucidated. Prior research also demonstrates that tau regulates N-methyl-D-aspartate (NMDA) receptor function by targeting Fyn Tyrosine Kinase to the post-synaptic density, where it phosphorylates NMDA receptors (26). Tau also has been shown to contribute to the stability of α-amino-3-hydroxy-5-methyl-4-isoxazolepropionic acid (AMPA) receptors through its interaction with the ATPase NSF (25).

The activity-regulated cytoskeleton-associated protein (Arc) is an immediate early gene that regulates diverse forms of synaptic plasticity, memory, and learning (27-31). One mechanism through which Arc regulates synaptic plasticity is by promoting the endocytosis of AMPA receptors through interactions with members of the endocytic machinery; endophilin-2/3, dynamin 2 and AP-2 that dominantly depends on its N-terminal region referred to as the endophilin binding (EB) domain (30,32,33).

Arc is upregulated during learning in the hippocampus (34) and is rapidly turned over, mainly through its ubiquitination and degradation by the ubiquitin proteasome system (UPS) (35-39). This type of posttranslational destabilization has been identified as a key mechanism for regulating group 1 metabotropic glutamate receptor-mediated long-term depression (mGluR-LTD) and spatial reversal learning (40). Arc is also removed by the autophagic lysosomal pathway (41) and can be degraded by noncanonical neuronal membrane associated proteasomes (42,43). Additionally, Arc undergoes several posttranslational modifications including, but not limited to, phosphorylation (44), sumoylation (45-47), palmitoylation (48), and acetylation (49).

Several studies have examined a role for Arc in AD pathology, mainly through examining the relationship between Arc and β-amyloid (50-54). However, there is a gap in understanding the relationship between Arc and tau. Given the role of Arc as a key regulator of synaptic plasticity and the recent implications of tau in regulating synaptic plasticity (55-57), we set out to determine if Arc might be affected by tau pathology. Here we show that endogenous tau has a physiological role in regulating Arc, where Arc levels are increased in the hippocampus of Tau KO mice and in dendrites of Tau KO primary hippocampal neurons. Conversely, overexpression of WT-tau but not P301L-tau led to reduced Arc stability. Tau-induced Arc reduction was found to be proteasome-dependent. Surprisingly, tau-dependent Arc degradation was not associated with increased Arc ubiquitination, lysosomal degradation or other known Arc posttranslational modifications that included phosphorylation, acetylation, or sumoylation. However, tau-dependent degradation did require the EB domain of Arc. Tau-induced alterations in Arc was selective to primary hippocampal dendrites and was associated with increased surface GluA1-containing AMPA receptors in dendrites and the soma. Our findings highlight a unique role of WT-tau in spatially and noncanonically regulating Arc removal, with hints of Arc endocytic targeting in regulating synaptic function.

## Results

Several studies show a role for tau in regulating synaptic plasticity. Arc levels are dysregulated in post-mortem brains from AD patients and in β-amyloid mouse models (54). We sought to investigate the potential relationship between Arc and tau, a hallmark of AD pathology, and whether Arc might be mediating some of the functions of tau at the synapse. We studied Arc protein in Tau KO mice, which lack the *Mapt* gene that encodes for Tau. Knock-out of *Mapt* was confirmed by genotyping and the absence of Tau protein (Supplemental Fig. 1A-B). We next compared Arc in hippocampi harvested from 3-month-old Tau KO mice and their WT littermates. Notably, Arc was significantly higher in total hippocampal lysates of Tau KO mice compared to WT (Fig. 1A; Arc, unpaired t-test t = 2.42, df = 12, p = 0.032). We asked if this increase was specific to a neuronal compartment. We biochemically fractionated extracts from hippocampus using serial centrifugations to isolate select subcellular fractions (Fig.1B). To demonstrate the effectiveness of our fractionation method, we analyzed GluA1 and the postsynaptic density protein (PSD-95) in isolated fractions. As expected, there was an increase in GluA1 and PSD-95 in the crude synaptosomal fraction (P2) and the lysed synaptosomal membrane fraction (P3) compared to the cytosolic fraction (S2) demonstrating successful fractionation (Supplemental Fig. 1C). Arc was significantly elevated in the P2 but not the S2 fraction, although there was a strong trend towards an upregulation of Arc in the S2 fraction (Fig. 1C, S2 t = 2.061, df = 11, p = 0.0638; Fig. 1D; Arc in P2, unpaired t-test t = 2.217, df = 12, p = 0.0467). No significant differences in Arc were found in the P3 and the synaptic vesicle (S3) fractions (Fig. 1E; Arc in S3, unpaired t-test t = 0.9787, df = 12, p = 0.347; Fig. 1F; Arc in P3, unpaired t-test t = 1.005, df = 12, p = 0.3349). Differences in GluA1 were not observed in any of the fractions from Tau KO mice.

**Figure 1:**
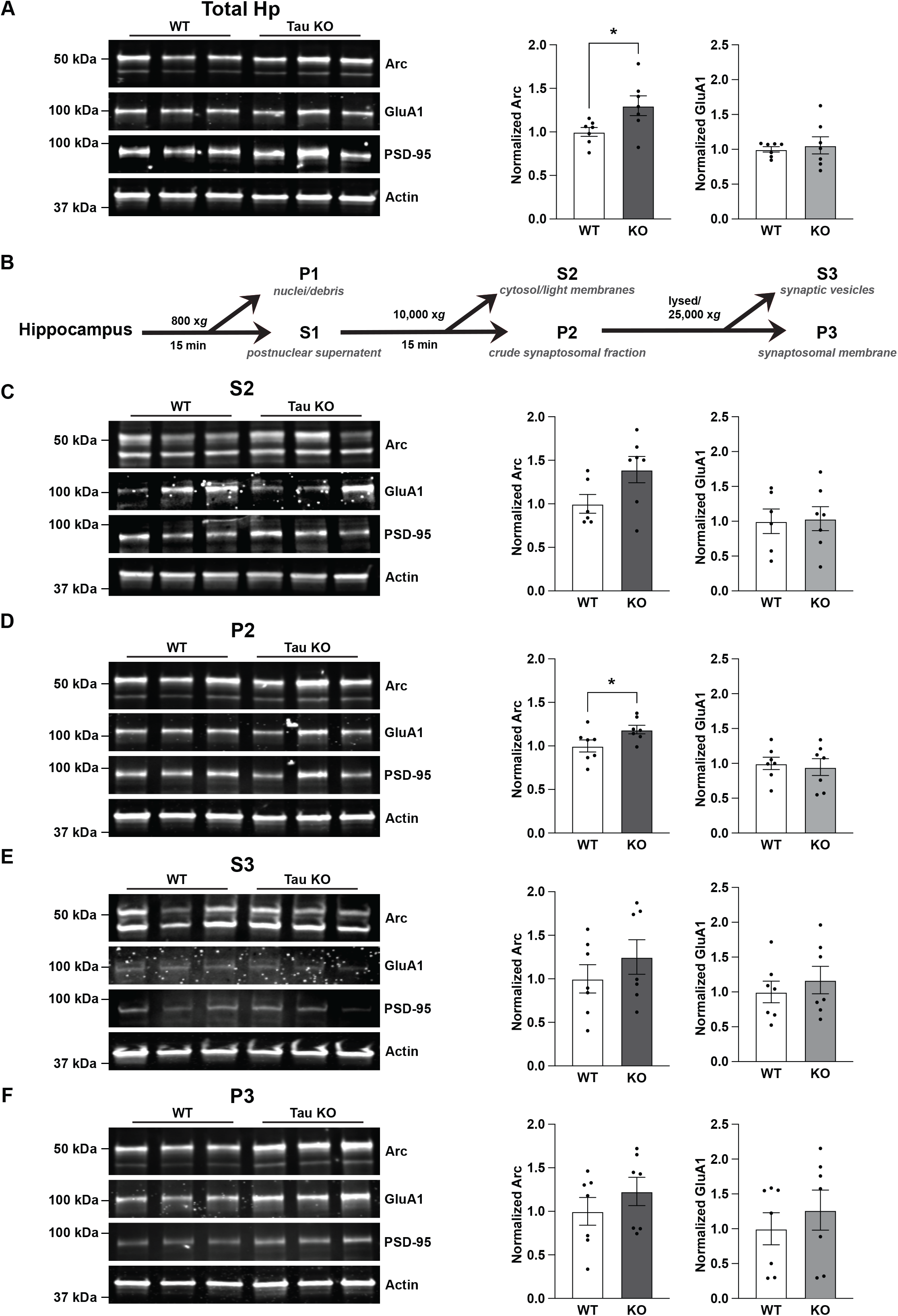
Arc is increased in the hippocampus of Tau KO mice. ***A,*** Representative western blots showing Arc, GluA1, PSD-95 and actin in total hippocampal (Hp) lysates. Hippocampi were harvested from 3 month-old WT and Tau KO littermates from both sexes. Arc was significantly higher in total Hp lysates. Unpaired t-test, t = 2.42, df = 12, p = 0.032. ***B,*** Biochemical fractionation scheme. ***C,*** Representative western blots showing Arc, GluA1, PSD-95 and actin in cytosol and light membrane fraction (S2). ***D,*** Representative western blots showing Arc, GluA1, PSD-95 in the crude synaptosomal fraction (P2). Arc was significantly higher in P2 fraction. Unpaired t-test, t = 2.217, df = 12, p = 0.0467. ***E,*** Representative western blots showing Arc, GluA1, PSD-95 and actin in cytosol in the synaptic vesicle fraction (S3). ***F,*** Representative western blots showing Arc, GluA1, PSD-95 and actin in the lysed synaptosomal membrane fraction (P3). No differences were found in GluA1 levels within any of the fractions. N = 6 animals per genotype, balanced for sex.

Since the increase in Arc was specific to the crude synaptosomal fraction, we investigated the spatial regulation of Arc by tau in primary hippocampal neuron cultures from WT and Tau KO littermates. tdTomato was used as a cell fill to outline neuronal morphology (Fig. 2A). Consistent with our hippocampal subcellular fractionation findings, Arc was selectively upregulated in dendrites and not the soma (Fig. 2B; unpaired t-test for Arc in dendrites, t = 2.517, df = 29, p = 0.0176; unpaired t-test for Arc in soma, t = 0.677, df = 29, p = 0.504). These findings suggest that tau plays a physiological role in regulating Arc selectively in dendrites.

**Figure 2:**
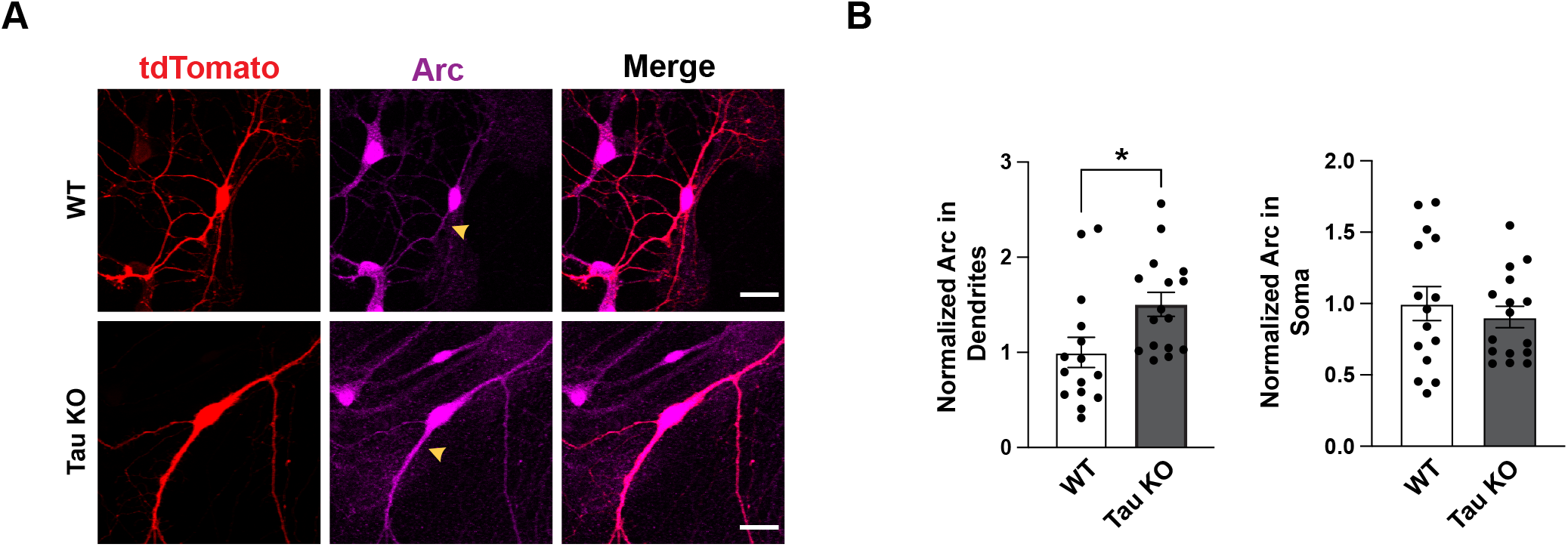
Arc is selectively upregulated in dendrites of Tau KO mice. ***A,*** Representative images of primary hippocampal neurons from WT and Tau KO littermates transfected at DIV 9 with tdTomato to outline neuron morphology and fixed at DIV 10. Scale bar = 20 μm ***B,*** Quantification of Arc showing significantly higher levels in dendrites but not the soma. Unpaired t-test for Arc in dendrites, t = 2.517, df = 29, p = 0.0176; unpaired t-test for Arc in soma, t = 0.677, df = 29, p = 0.504. n = 15-16 neurons from 2 independent biological replicates.

We asked if high levels of tau, similar to those observed in tauopathies would also affect Arc in neurons. GFP-tagged tau (GFP-tau) was overexpressed in WT primary hippocampal neurons and endogenous Arc levels were quantified. tdTomato was used as a cell fill to outline neuronal morphology (Fig. 3A). Consistent with the increase in Arc selectively in dendrites of Tau KO mice, we found that Arc protein was selectively decreased in dendrites upon GFP-tau overexpression. We also investigated the effect of GFP-P301L tau overexpression on Arc. P301L-tau is a missense single-point mutation located on the R2 MTBD that substitutes Proline for Leucine and has been commonly used to model AD pathology (17). Unlike GFP-tau, overexpression of GFP-P301L tau had no effect on Arc in soma or dendrites (Fig. 3B; One-way ANOVA in dendrites F(2,55) = 5.885, p = 0.0048, Tukey’s post-hoc GFP vs. GFP-Tau p = 0.0023; one-way ANOVA in soma F (2,62) = 0.62, p = 0.54).

**Figure 3:**
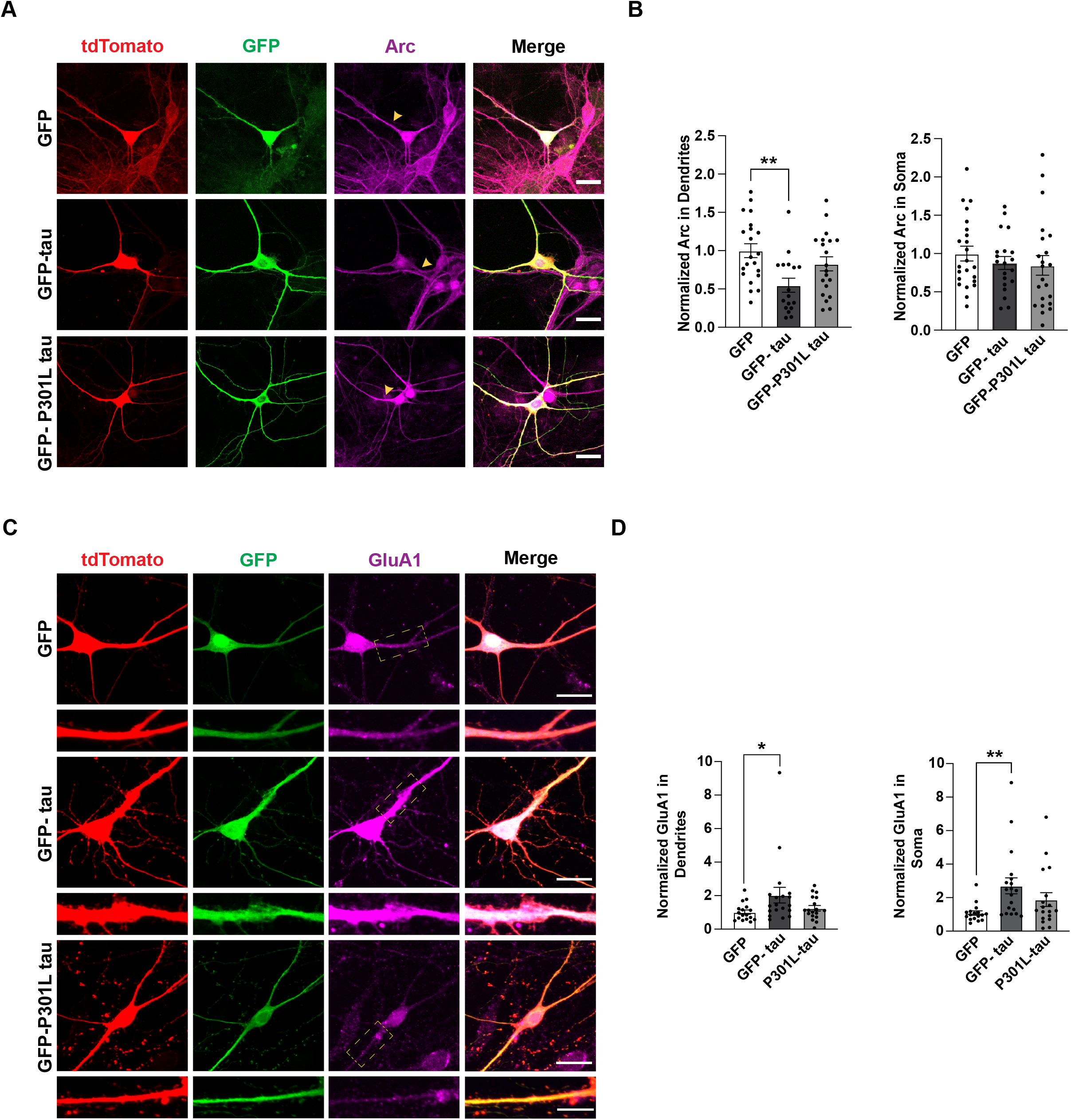
GFP-tau but not GFP-P301L tau selectively reduces Arc in dendrites and correspondingly increases surface GluA1 in primary hippocampal neurons. ***A***, Primary hippocampal neurons transfected with GFP, GFP-tau or GFP-P301L tau. Tdtomato was used to outline neuron morphology. Scale bar represents 20 μm. ***B,*** Quantification of Arc protein (magenta) in soma and dendrites showing a decrease in Arc selectively in dendrites with GFP-tau but not with GFP-P301L. One-way ANOVA in dendrites F(2,55) = 5.885, p = 0.0048, Tukey’s post-hoc GFP vs. GFP-Tau p = 0.0023; one-way ANOVA in soma F (2,62) = 0.62, p = 0.54. n = 16-21 neurons from 3 independent biological replicates. ***C,*** Primary hippocampal neurons overexpressing GFP, GFP-tau and GFP-P301L tau. tdTomato was used to outline neuron morphology. Neurons were fixed and then immunostained with an anti-GluA1 antibody. Scale bar represents 20 μm. Scale bar for selected dendrites represents 10 μm. ***D***, Quantification of surface GluA1 (magenta) in soma and apical dendrites. Kruskal-Wallis test for dendrites, p = 0.0458; Kruskal-Wallis test for soma, p = 0.0034. n = 17-19 neurons from 3 independent biological replicates.

Given the role of Arc in regulating AMPA receptor trafficking (32), we hypothesized that the tau-mediated decrease of Arc in dendrites may result in decreased AMPA receptor endocytosis and consequently lead to an increase in surface AMPA receptor levels. To test this, we overexpressed GFP-tau and GFP-P301L tau in primary hippocampal neurons and quantified surface GluA1-containing AMPA receptors (Fig. 3C). GluA1 staining in neurons overexpressing GFP-tau had a smoother appearance, unlike the punctate distribution in neurons overexpressing GFP and GFP-P301L tau. As expected, GFP-tau overexpression led to an increase in surface GluA1; however, this effect was not selective to dendrites, as overexpression also led to an increase of surface GluA1 in the soma (Fig. 4D, Kruskal-Wallis test for dendrites, p = 0.0458, Dunn’s multiple comparisons test, GFP vs GFP-tau p = 0.0415; Kruskal-Wallis test for soma, p = 0.0034, Dunn’s multiple comparisons test, GFP vs GFP-tau p = 0.0023). Since Arc may play a role in regulating dendritic spine density (58), we quantified the number of dendritic spines in each of the conditions. However, we found no significant differences in spine density compared to GFP alone (Supplemental Fig. 2, Ordinary one-way ANOVA F (2, 50) = 0.1543, p = 0.8574). These findings suggest that WT but not GFP-P301L tau overexpression decreases Arc and subsequently increases surface expression of GluA1-containing AMPA receptors.

**Figure 4:**
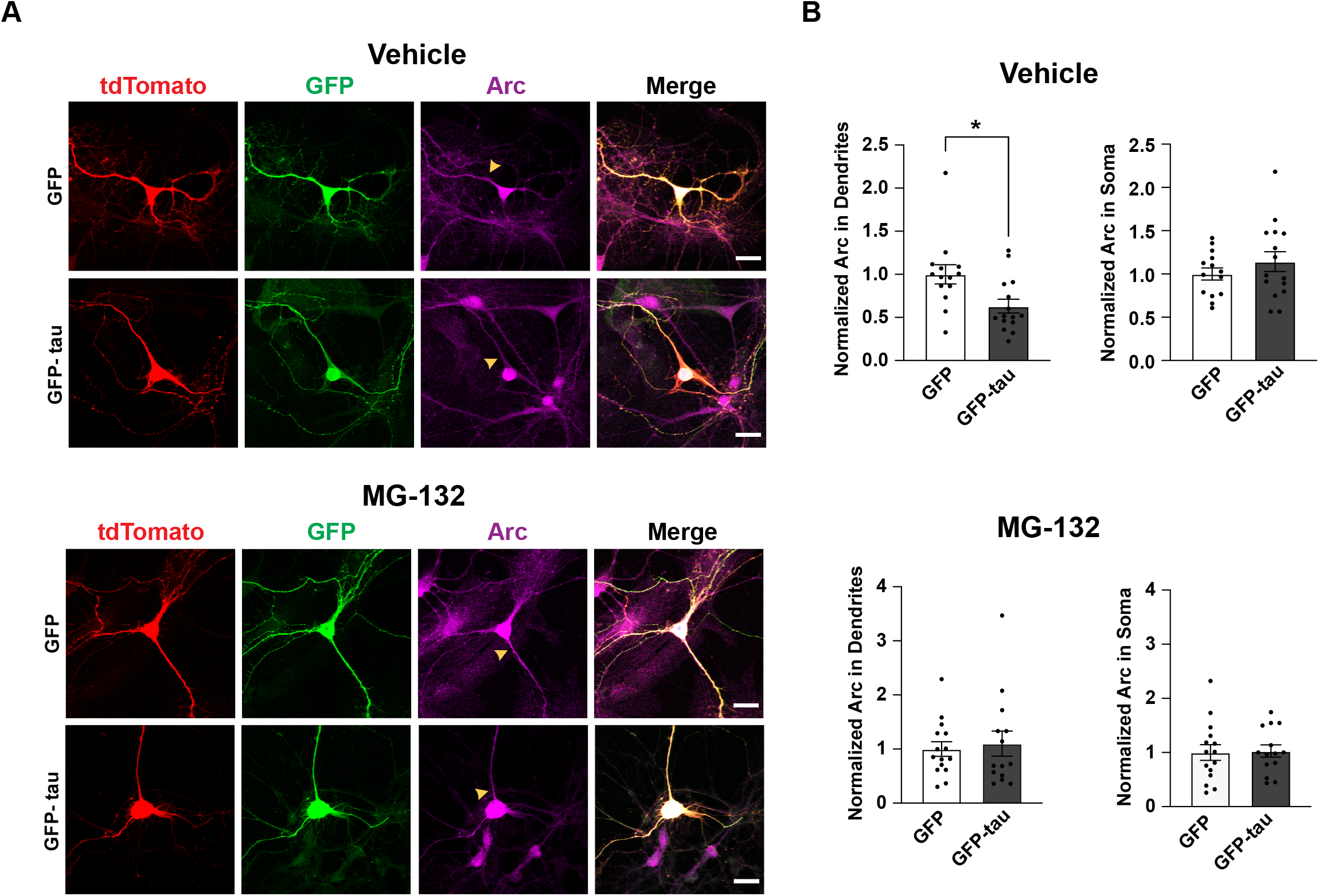
Tau modulation of dendritic Arc in primary hippocampal neurons is proteasome-dependent. ***A,*** Primary hippocampal neurons overexpressing GFP or GFP-tau. tdTomato was used to outline neuron morphology. Cells treated with either Vehicle (DMSO) or MG-132 (10 μM) for 4 h. Scale bar represents 20 μm. ***B,*** Quantification of Arc protein (magenta) in soma and apical dendrites showing a decrease in Arc selectively in dendrites in cultures treated with Vehicle but not in cultures treated with MG-132. Unpaired t-test DMSO Control in dendrites t=2.72, df = 27, p = 0.011; Unpaired t-test MG-132 in dendrites t = 0.3814, df = 27 p = 0.706; Unpaired t-test DMSO in soma t = 1.044, df = 27, p = 0.306; Unpaired t-test MG-132 in soma t = 0.16, df = 27, p = 0.874. n = 14-15 neurons from 2 independent biological replicates.

We sought to determine the mechanism through which tau regulates Arc expression. One possibility is that tau could be regulating Arc stability at the posttranslational level. Ubiquitination of Arc is a modification that facilitates its degradation by the proteasome, which is a major pathway for its posttranslational removal (35,36,39). Therefore, we asked whether the reduction of Arc by tau was dependent on proteasome activity. To determine if tau-dependent modulation of Arc was proteasome-dependent, primary hippocampal neurons overexpressing GFP-tau were treated with the proteasome inhibitor MG-132 and Arc was quantified (Fig. 4A). In Vehicle treated neurons, Arc was significantly decreased with GFP-tau overexpression but not in MG-132-treated neurons (Fig. 4B; Unpaired t-test DMSO Control in dendrites t = 2.72, df = 27, p = 0.011; Unpaired t-test MG-132 in dendrites t = 0.3814, df = 27 p = 0.706; Unpaired t-test DMSO in soma t = 1.044, df = 27, p = 0.306; Unpaired t-test MG-132 in soma t = 0.16, df = 27, p = 0.874). Our findings support the notion that tau modulation of Arc is selective for dendrites and is proteasome-dependent.

We next turned to HEK293 cells, which are more amenable to performing biochemical studies to further elucidate the possible multitude of mechanisms through which WT-tau regulates Arc. Cells overexpressing myc-tagged Arc and increasing concentrations of GFP-tau were treated with the proteasome inhibitor MG-132 (Fig. 5A). Similar to the observations with endogenous Arc in primary hippocampal neurons, myc-Arc was reduced upon increasing concentrations of GFP-tau but this effect was not observed in cells treated with MG-132 (Fig. 5B; one-way ANOVA for DMSO control, F (5,12) = 4.03, p = 0.022, Tukey’s post-hoc test 0 vs. 1 μg: p= 0.046; one-way ANOVA for MG-132, F (5,12) = 1.9, p = 0.15). We also overexpressed myc-tagged Arc with increasing concentrations of GFP-P301L tau and found no significant difference in myc-Arc with increasing GFP-P301L tau. (Supplemental figure 3, One-way ANOVA, F (5,18) = 0.3018, p = 0.3).

**Figure 5:**
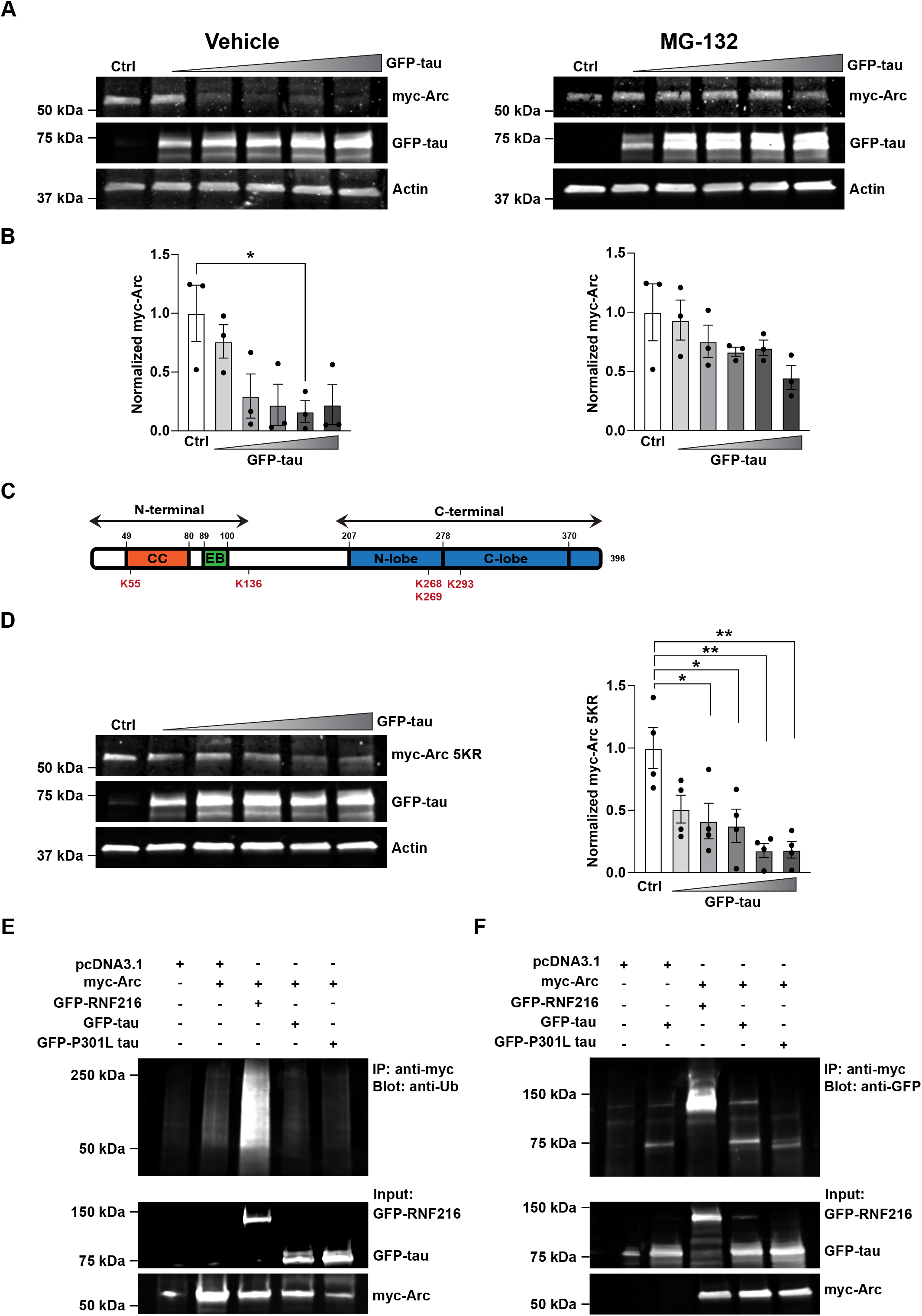
Tau modulation of Arc in HEK293 cells requires proteasome activity but does not increase Arc ubiquitination. ***A,*** Following transfection, HEK293 cells were treated with either Vehicle (DMSO) or MG-132 (10 mM) for 4 hours. *Left, r*epresentative western blots showing myc-Arc with increasing concentrations of GFP-tau in DMSO treated HEK293 cells (0, 0.25, 0.5, 0.75, 1 and 1.5 μg). Actin was used as a loading control. *Right*, *r*epresentative western blots showing myc-Arc with increasing concentrations of GFP-tau in DMSO treated HEK293 cells (0, 0.25, 0.5, 0.75, 1 and 1.5 μg). Actin was used as a loading control. pcDNA3.1 was used as a DNA filler to keep the amount of transfected DNA between titration conditions identical. ***B,*** Quantification of myc-Arc normalized to actin showing a significant decrease in Arc with increasing Tau in vehicle-treated cells (left), but not in the MG-132-treated cells (right). One-way ANOVA for DMSO control, F (5,12) = 4.03, p = 0.022; one-way ANOVA for MG-132, F (5,12) = 1.9, p = 0.15. n = 3. ***C,*** Schematic showing the structure of Arc, highlighting the coiled-coil (CC) domain and the endophilin-binding (EB) domain on the N-terminus, and the N- and C-lobe on the C-terminus. The locations of the lysines (K55,136, 268, 269 and 293) targeted for ubiquitination by RNF216 and UBE3A are shown. ***D,*** *Left,* Representative western blots showing myc-Arc5KR (K55,136, 268, 269 and 293 mutated to Arginine) with increasing concentrations of GFP-tau in HEK293 cells (0, 0.25, 0.5, 0.75, 1 and 1.5 μg). Actin was used as a loading control. *Right*, Quantification of myc-Arc5KR levels normalized to actin showing a significant decrease in Arc with increasing tau. One-way ANOVA F = (5,18) = 6.4, p = 0.0014. pcDNA3.1 was used as a DNA filler to keep the amount of transfected DNA between titration conditions identical. n = 4. ***E,*** *Top,* Ubiquitin assay showing no enhancement in Arc ubiquitination when co-expressed with Tau or P301L tau. RNF216 was used as a positive control. *Bottom*, Input showing expression of myc-Arc, GFP-RNF216, GFP-tau and GFP-P301L tau. ***F,*** Coimmunoprecipitation assay showing pulldown of myc-Arc with anti-myc antibody and immunoblotting with an anti-GFP antibody. GFP-RNF216 was used as a positive control.

Studies have shown that Arc can be ubiquitinated by the E3 ligases RNF216, UBE3A and CHIP, which induces its rapid degradation by the proteasome (35,36,38). Since protein ubiquitination dominantly occurs on lysine residues, we tested reported Arc ubiquitination sites for RNF216 and UBE3A on lysines which were mutated to arginine to prevent ubiquitination (K55R/K136R/K268R/K269R/K293R, myc-Arc5KR) (35,36) (Fig. 5C). Surprisingly, co-transfection of myc-Arc5KR with increasing amounts of GFP-tau still led to a reduction in Arc that was similar in magnitude to myc-Arc WT (Fig. 5D; one-way ANOVA F = (5,18) = 6.4, p = 0.0014, Tukey’s post-hoc 0 vs. 0.5 μg = 0.02, 0 vs 0.75 μg p = 0.0175, 0 vs. 1 μg p = 0.0015, 0 vs. 1.5 μg p = 0.0016). These findings suggested that preventing Arc ubiquitination at these sites does not interfere with tau-mediated Arc reduction. To determine if the tau-mediated decreases in Arc involved ubiquitination, we measured myc-Arc ubiquitination in the presence of GFP-tau or GFP-P301L tau after treatment with MG-132 to trap Arc ubiquitinated products. While RNF216 robustly increased myc-Arc ubiquitination, this was not observed upon GFP-tau or GFP-P301L tau overexpression (Fig. 5E). To test if Arc could interact with WT-tau, we performed co-immunoprecipitation assays with WT or P301L-tau. While RNF216 efficiently co-immunoprecipitated with Arc, interaction with WT or P301L-tau was not different compared to background binding control (Fig. 5F). Taken together, these experiments suggest that even though tau modulation of Arc is proteasome dependent, it does not appear to be through a direct interaction with Arc or through enhancing Arc ubiquitination by RNF216 or UBE3A, which indicates that tau might be utilizing noncanonical methods of Arc removal by the proteasome.

One alternative possibility to our MG-132 results could be related to the ability of MG-132 to block late phases of lysosomal degradation (59). Given that Arc is also removed by the autophagy lysosome system in neurons (41), we asked if tau decreases Arc by lysosome-dependent degradation. We treated cells overexpressing myc-Arc and GFP-tau with the lysosome inhibitors Leupeptin and Ammonium Chloride for 6 h before harvest (Supplemental Fig. 4A). We confirmed inhibition of lysosomal activity by blotting for markers of autophagic flux – MAP1LC3B (hereafter LC3) and p62/SQSTM1 (60). There was a significant increase in the autophagosome bound form of LC3 – LC3-II (Supplemental Fig. 4B; unpaired t-test, t = 12.52, df = 22, p < 0.0001) and the autophagosome substrate p62/SQSTM1 (unpaired t-test, t = 10.54, df = 22, p < 0.0001) in cells treated with lysosome inhibitors as compared to vehicle treated cells. Surprisingly, we did not observe the expected increase in Arc protein (Unpaired t-test, t = 0.7087, df = 10, p = 0.4947). Nevertheless, Arc was decreased upon GFP-tau overexpression in both vehicle and inhibitor-treated conditions (Supplemental Fig. 4C; Unpaired t-test for vehicle control, t = 10.6, df = 10, p < 0.0001; Unpaired t-test for inhibitors, t = 5.585, df = 10, p = 0.0002). These findings demonstrate that tau modulation of Arc is not lysosome dependent.

Given the ability of overexpressed tau to form insoluble aggregates (61,62), we next asked if tau may be precipitating Arc into insoluble aggregates that could not be extracted in the RIPA-soluble fraction. We extracted proteins in the RIPA-insoluble fraction using formic acid, which has been successfully used to extract tau aggregates (63,64). While GFP-tau was detected in the insoluble fraction, Arc was not (Supplemental Figure 4D), indicating that tau-dependent decreases in Arc are not due to its precipitation into insoluble aggregates.

One recent study suggested that Arc phosphorylation by GSK3α/β can enhance Arc removal by the proteasome (44). We treated cells co-expressing myc-Arc and GFP-tau with the GSK3α/β inhibitor CHIR 98014 (CH98) for 4 h prior to harvest (Fig. 6A). myc-Arc decreased with GFP-tau overexpression in both vehicle and the CH98-treated condition (Fig. 6A; Unpaired t-test for vehicle control, t = 3.450, df = 18, p = 0.0029; unpaired t-test for CH98, t = 3.417, df = 18, p = 0.0031). It was also reported that Arc is phosphorylated on S170, T175, T368, and T380 by GSK3α/β (44) (Fig. 6B). We mutated these phosphorylation sites to generate myc-Arc S170A/T175A, myc-Arc T368, and myc-Arc T380A. However, upon overexpression with GFP-tau (Fig. 6, C-E), all 3 of these myc-Arc phosphorylation mutants were still decreased (Fig. 6, C-E; Unpaired t-test for Arc S170A/T175A, t = 4.913, df = 16, p = 0.0002; Unpaired t-test for Arc T368A, t = 8.714, df = 4, p = 0.001; Unpaired t-test for Arc T380A, t = 11.59, df = 4, p = 0.0003). Together, these experiments demonstrate that tau modulation of Arc is not mediated through GSK3α/β activity or GSK3α/β-dependent Arc phosphorylation.

**Figure 6:**
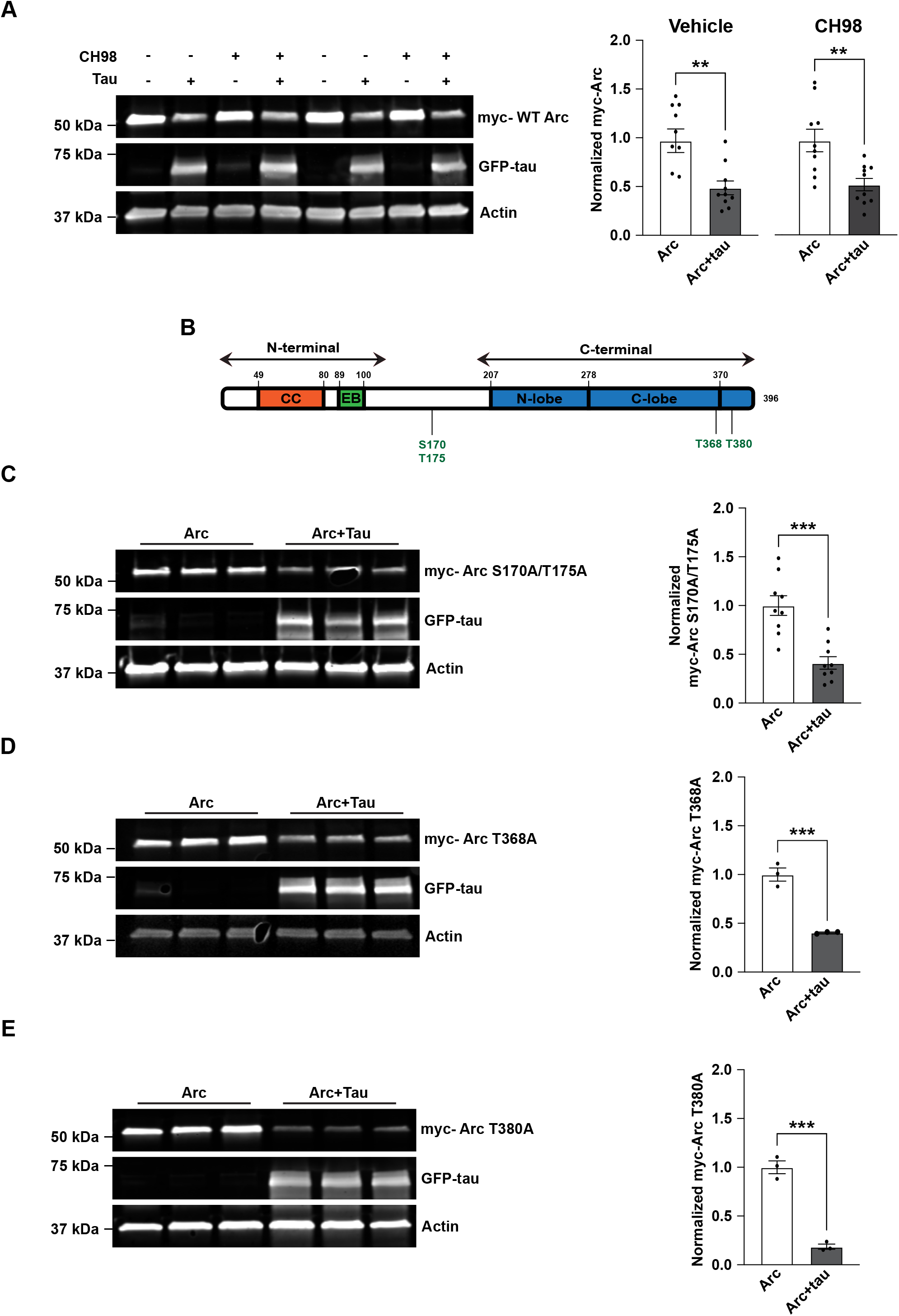
Tau-induced Arc removal does not depend on GSK3α/β Arc phosphorylation. ***A,*** *Left,* Representative western blots showing myc-Arc expressed alone or with GFP-tau. Cells were treated with Vehicle (Water) or CH98 (1-2 μM) for 4 h. Actin was used as a loading control. Right, Quantification of myc-Arc showing a significant decrease with co-expression of GFP-tau after treatment with Vehicle (unpaired t-test, t = 3.450, df = 18, p = 0.0029) or CH98 (unpaired t-test, t = 3.417, df = 18, p = 0.0031). n = 9 ***B,*** Schematic showing the structure of Arc with the location of mapped GSK3α/β phosphorylation sites S170, T175, T368 and T380. ***C-E,*** *Left,* Representative western blots showing myc-Arc S170A/T175A, myc-Arc T368A, or myc-Arc T380A with Arc phosphorylation sites mutated to Alanine expressed alone or with GFP-tau. Actin was used as a loading control. *Right,* Quantification of myc-Arc S170A/T175A, myc-Arc T368A, or myc-Arc T380A showing a significant decrease when co-expressed with Tau. Unpaired t-test for Arc S170A/T175A, t = 4.913, df = 16, p = 0.0002; Unpaired t-test for Arc T368A, t = 8.714, df = 4, p = 0.001; Unpaired t-test for Arc T380A, t = 11.59, df = 4, p = 0.0003. n = 9 for S170A/T175A, n = 3 for T368A and T380A.

The lack of effects of ubiquitin and phosphorylation modifying sites to mediate Arc removal by tau prompted us to evaluate larger regions of Arc that might be necessary for tau-dependent reductions. We used myc-Arc constructs that lack specific domains of Arc; myc-Arc ΔC-terminal (lacking the C-terminal domain), myc-Arc ΔCC (lacking the coiled-coil motif on the N-terminal domain), and myc-Arc ΔEB (lacking the EB domain on the N-terminus) (36) (Fig. 7A). We found that all the tested myc-Arc constructs were significantly decreased with tau except for the myc-Arc ΔEB (Fig. 7B; Unpaired t-test for WT Arc t = 8.857, df = 10, p < 0.0001; Unpaired t-test for Arc ΔC-terminal t=6.471, df = 10, p < 0.0001; Unpaired t-test for Arc ΔCC t = 7.076, df = 10, p < 0.0001; Unpaired t-test Arc ΔEB t = 0.2956, df =1 0, p = 0.774). Cumulatively, these findings suggest that the EB domain of Arc is essential for its reduction by tau.

**Figure 7:**
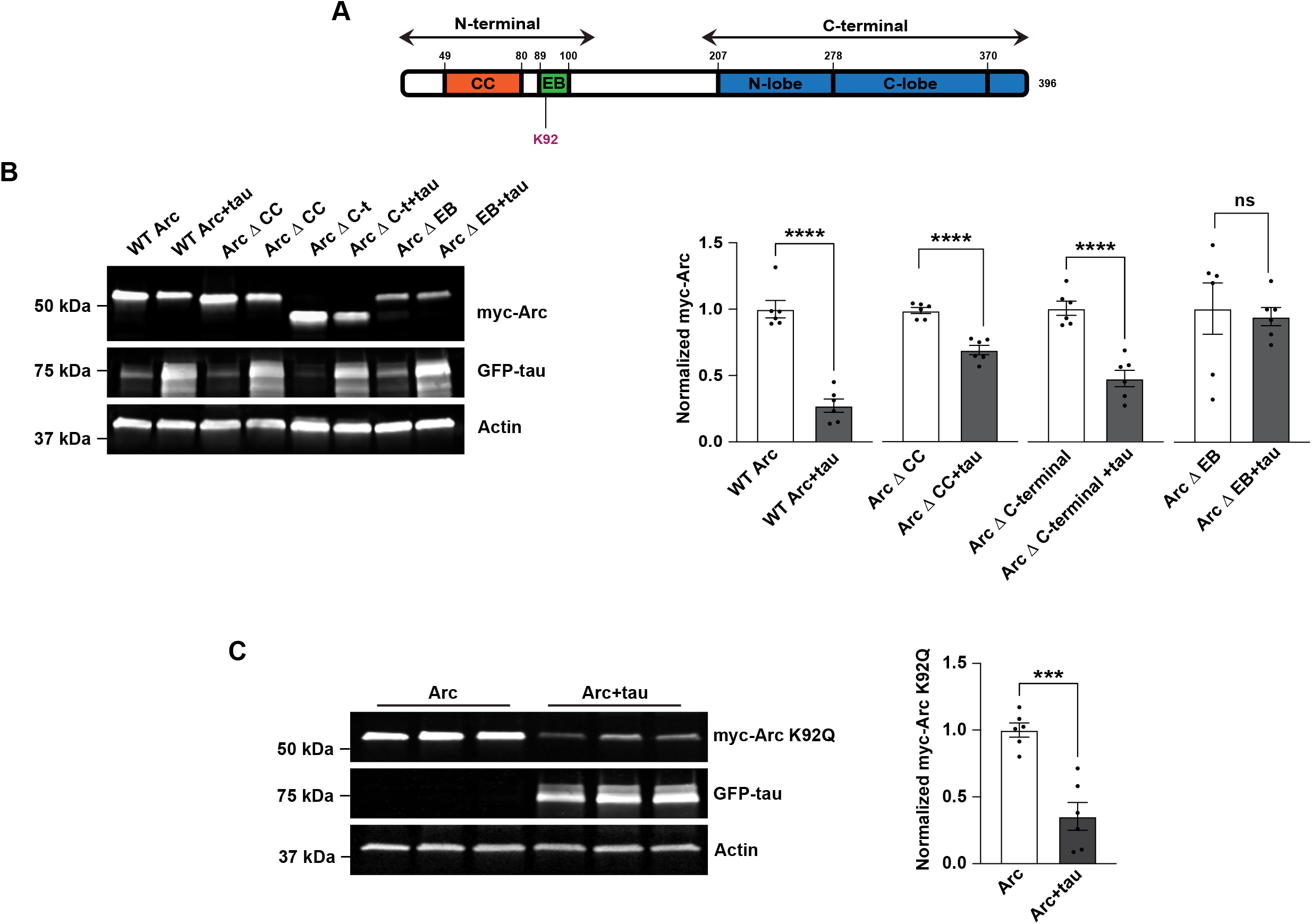
The Endophilin-binding domain of Arc is essential for Arc modulation by tau. ***A,*** Schematic showing the structure of Arc, highlighting the coiled-coil (CC) domain and the endophilin-binding (EB) domain on the N-terminus, and the N- and C-lobe on the C-terminus. The location of the K92 acetylation site is shown. ***B,*** *Left,* Representative western blot showing WT myc-Arc, myc-Arc ΔC-terminal (lacking the C-terminal domain), myc-Arc ΔCC (lacking the coiled-coil motif on the N-terminal domain), and myc-Arc ΔEB (lacking the endophilin-binding domain on the N-terminus) expressed alone or with GFP-tau. Actin was used as loading control. *Right,* Quantification of WT myc-Arc, myc-Arc ΔC-terminal, myc-Arc ΔCC, and myc-Arc ΔEB. Only Arc ΔEB does not show a decrease with Tau overexpression (Unpaired t-test for WT Arc t = 8.857, df = 10, ****p < 0.0001; Unpaired t-test for Arc ΔC-terminal t = 6.471, df =1 0, ****p < 0.0001; Unpaired t-test for Arc ΔCC t = 7.076, df = 10, ****p < 0.0001; Unpaired t-test Arc ΔEB t = 0.2956, df = 10, p = 0.774). n = 6. ***C,*** *Left,* Representative western blots showing myc-Arc K92Q with the acetylation site K92 mutated to glutamine expressed alone or with GFP-tau. Actin was used as a loading control. *Right,* Quantification of myc-Arc K92Q showing a significant decrease when co-expressed with Tau. Unpaired t-test t = 5.54, df = 10, p = 0.0002. n = 6.

The Arc EB domain is an 11 amino acid sequence that is important for targeting Arc to endosomes (32). Recently, Arc was found to be acetylated at K24, K33, K55 and K92 which increases its stability (49). Of these sites, K92 falls within the EB domain (89-100) (Fig. 7A). We hypothesized that tau could be decreasing Arc stability by interfering with its acetylation at K92. To test this hypothesis, we created the acetyl-mimetic myc-Arc K92Q, which increases the stability of Arc (49). However, the myc-Arc K29Q mutant was still reduced with GFP-tau overexpression suggesting that tau does not modulate Arc through interfering with its acetylation at K92 (Fig. 7C; Unpaired t-test t = 5.54, df = 10, p = 0.0002). Cumulatively, these findings suggest that tau regulation of Arc is not occurring through canonical degradation pathways and known posttranslational modifications.

## Discussion

Here we show that the regulation of Arc by tau is physiological, and not limited to conditions of tau overexpression. We found that Arc is upregulated in hippocampal lysates and in the crude synaptosomal subcellular fraction of Tau KO mice. Arc is also selectively upregulated in dendrites of Tau KO primary hippocampal neurons. Overexpression of tau led to the opposite effect; reducing Arc selectively in neuronal dendrites. Tau-dependent decreases in Arc in dendrites is proteasome-dependent, suggesting that tau regulates posttranslational Arc stability. The decrease of Arc with tau overexpression was also associated with an increase in surface expression of GluA1 in both soma and dendrites. In attempting to decipher the mechanism through which tau regulates Arc, we tested the role of numerous Arc post-translational modifications in HEK293 cells that could be responsible for regulating tau-dependent Arc turnover. Despite dependence on the proteasome, tau regulation of Arc was independent of Arc ubiquitination, phosphorylation, acetylation and lysosomal degradation, suggesting that tau regulation of Arc occurs through non-canonical pathways. However, we show that the endophilin-binding domain of Arc is necessary for its modulation by tau, suggesting a role for Arc engagement with the endocytic machinery for tau-induced instability.

Several studies have shown a role for Arc in AD, mainly through links to amyloid-beta (Aβ). For example, both increases and decreases in Arc in the hippocampus and cortex were reported in several amyloid precursor protein (APP) mouse models (50-53,65-67) and it has been suggested that these changes occur in an age-dependent manner (68,69). Arc levels are also increased in the medial prefrontal cortex of patients with AD (54). On the other hand, a mechanistic relationship between Arc and tau has been relatively understudied. While experience-driven Arc responses were found to be disrupted in the vicinity of plaques in the APP/PS1 model, where neurons in the vicinity of amyloid plaques were less likely to respond, no similar effect was observed in the vicinity of tau NFTs in the Tg4510 mouse overexpressing P301L tau (70,71). A recent study showed that tau elevated Arc1 in a drosophila AD model overexpressing R406W tau, a mutant linked to Frontotemporal dementia, demonstrating a role for Arc1 in neurodegeneration (72). While the conflicting results make it difficult to define a clear role for Arc in AD pathology, this can be attributed to differences in species, the disease models used, the stage of disease development, and the tissue type. Differences in levels of excitability between networks and brain areas can also explain some of the contradictions as Arc levels increase rapidly in excited synapses and upon exposure to learning experiences followed by its removal to return to its basal levels (36,68,73-75). We show that tau modulation of Arc is proteasome dependent. A major pathway for removal of Arc is through the UPS (35-39). However, creating mutations of characterized Arc ubiquitination sites targeted by the E3 ligases RNF216 and UBE3A (Arc5KR) did not block Arc degradation (35,36). Although we did not test for a role of CHIP, an E3 ligase that regulates the ubiquitination of both Arc and tau (38,76), the possibility remains unlikely as we could not detect an enhancement of Arc ubiquitination upon tau overexpression. Moreover, tau modulation of Arc was not dependent on Arc phosphorylation or lysosomal degradation. In light of these findings, three alternative mechanisms can still be hypothesized. First, recent studies have found that Arc can be assembled into viral-like capsids and released into the extracellular space (77-79). It is possible that tau overexpression may result in the extracellular release of Arc capsids. Second, Arc is a substrate of the neuronal membrane proteasome (NMP), which is proteasome inhibitor sensitive and utilizes a ubiquitin-independent mechanism for degradation of ribosome-associated nascent Arc (42,43). However, HEK293 cells do not express the NMP yet we found that tau overexpression was still able to reduce Arc in a proteasome-dependent manner. A final possibility is that Arc may be degraded by 20S uncapped proteasomes, which are MG-132 sensitive and can also function to degrade proteins independently of protein ubiquitination (80). Both neuronal and nonneuronal cells express 20S uncapped proteasomes at differing abundances (81,82).

We show that the EB domain mutant of Arc is required for modulation by tau. Structurally, Arc consists of two juxtaposed domains, a positively charged N-terminal domain (NTD) and a negatively charged C-terminal domain (CTD). The NTD has several peptide-binding sites, including the EB and two long helices possibly forming a coiled-coil (83,84). Arc mediates endocytosis of AMPA receptors through its interaction with Endophilin at the EB domain (32). Given our findings requiring the EB of Arc, we investigated the possibility that tau overexpression could be interfering with Arc stability by disrupting the acetylation of Arc at K92 located within this domain (49), but an acetyl mimetic Arc mutant at this site (K92Q) did not prevent the decrease in Arc. Moreover, Arc did not co-immunoprecipitate with tau even when blocking its degradation with the proteasome inhibitor MG-132, suggesting an indirect or transient interaction between these two proteins. Arc binds endophilin, dynamin, and AP-2 to mediate endocytosis of AMPA receptors (32,33). Recent findings show that tau (2N4R) has an extensive network of interactions with proteins that regulate endocytosis, including endophilin, AP-2 and dynamin (25,85). Interestingly, expression of human tau (0N4R) induces de novo assembly of microtubules which interferes with endocytosis through sequestration of dynamin (86). Although we are unable to define a detailed mechanism for the modulation of Arc by tau, our results suggest the involvement of the endocytotic machinery, engaged by both Arc (via EB domain) and tau, in this process. However, it is important to be careful with interpretation of data from a heterologous expression system such as HEK293 cells. While we show that the decrease in Arc by tau is proteasome-dependent in both neurons and HEK293 cells, the upstream mechanisms involved in targeting of endogenous Arc to the proteasome in primary hippocampal neurons could potentially be different from mechanisms of removal of artificially expressed myc-Arc in HEK 293 cells. This becomes particularly important in studies such as ours that evaluate alterations in levels of artificially expressed proteins. We controlled for the efficiency of transfection by keeping the amount of DNA transfected into the cell identical in all conditions by adding the filler plasmid pcDNA3, which may be a potential confound in some of our conclusions from experiments conducted solely in HEK293 cells.

Consistent with the observed decrease in Arc specifically in hippocampal dendrites upon tau overexpression, we found that WT tau overexpression also increases surface levels of GluA1 in the soma of primary hippocampal neurons and in dendrites. Since surface GluA1 was also increased in the soma despite no detected change in Arc in that region, we propose that tau overexpression might mistarget GluA1-containing AMPA receptors to the soma due to altered trafficking. The physiological interactions of dendritic tau with synaptic proteins that regulate postsynaptic receptor trafficking and synaptic plasticity have been described (23,25,55,87,88). However, we did not observe a change in GluA1 levels in subcellular fractions of Tau KO mouse brains. A study that investigated surface GluA1 levels in hippocampal neurons of Tau KO mice under different conditions showed no change in basal levels of surface GluA1 in neurites, but a decrease in total surface GluA1 compared to WT controls (25). Tau interacts with several proteins that regulate AMPA receptor trafficking profiles such as NSF and PICK1 (25,88), and tau knockout neurons show a rapid reduction in the number of GluA2 puncta after NMDA treatment (89). Additionally, tau plays a role in the postsynaptic targeting of the Src kinase Fyn, which regulates the activation of NMDA receptors (55), and consequently affects AMPA receptor trafficking (90). Thus, the observed results could be a net effect of the interaction of tau with several proteins. The sequestration of these proteins could lead to Arc instability. In contrast, we did not observe a change in surface GluA1 with P301L tau overexpression in the soma or dendrites of hippocampal neurons. Hoover et al. reported higher levels of P301L tau in postsynaptic density proteins isolated from rTg4510 mice overexpressing P301L tau compared to those isolated from rTg21221 mice overexpressing WT tau (0N4R). rTgP301L neurons showed lower spine GluA1 levels compared to neurons from rTgWT mice at DIV 21-35 (91). In our study, we did not find any changes in dendritic spine densities between WT and P301L tau 48 hours post transfection. These discrepancies may be due to acute versus chronic overexpression and thus our findings may model an earlier stage of tau pathology. Additionally, the insertion of the transgene *MAPT P301L* in the Tg4510 model disrupts the *fibroblast growth factor 14* (*Fgf14*) gene, which has been shown to contribute to the neurodegeneration observed in the rTgP301L mouse model, which may have inadvertently resulted in off target effects on surface GluA1 and spine densities (92,93).

While different tau isoforms and mutants are used interchangeably to model AD and other tauopathies, our study adds to the growing body of literature that emphasizes the differences in protein interactions between tau and its mutants, which would suggest that they have different roles in the cell as well as different contributions to disease pathogenesis (22,85,94-96). The six isoforms of tau are differentially expressed throughout development, with the ratio of 3R to 4R tau in the adult human brain roughly equal to one (8). Tau isoforms further have distinct biochemical properties such as different propensities for aggregation, with those containing 4R assembling 2.5 to 3 times faster compared to 3R isoforms (97). In AD, NFTs contain all six isoforms, while in other tauopathies, tangles may predominantly have 3R or 4R tau (98). On the other hand, the majority of *MAPT* mutations, including P301L, are associated with FTD. However, P301L as well as other *MAPT* mutations have been commonly used to model AD in vivo and in vitro despite their distinct physical properties (99). The genetically matched rT1 model overexpressing WT 0N4R human tau and rT2 model overexpressing P301L-tau also show differences in tau phosphorylation and stability at different developmental stages (100).

Evidence of involvement of Arc in AD pathology, and the role of Arc in regulating learning and memory which are severely disrupted in AD, raises the question to the potential of targeting Arc therapeutically to ameliorate some of these disruptions. Several drugs are known to alter Arc levels and function, including psychotropic drugs as well as other drugs that manipulate the proteasome and the autophagy-lysosome systems, most of which are well-studied and are already in use to treat other disorders (101). However, given the complexity of the role of Arc in AD, the desired effect of pharmacologically altering Arc remains unclear. More studies are needed to evaluate a well-defined role for Arc in AD that takes into consideration the role of Arc in regulating AD pathology such as Aβ, and the effect of tau pathology on regulating Arc. Notably, Arc is robustly induced with experiences that stimulate plasticity and is specifically targeted to stimulated synapses (102), and holistic approaches that are already in practice for AD management such as cognitive therapy, have shown evidence of substantial benefits for AD patients, and many of them can induce Arc in a non-pharmacological manner (103). Additionally, several drugs designed to reduce tau through immunotherapy are currently in clinical trials, which can possibly ameliorate the downstream effects of increased tau (104).

Our study identifies a new physiological role for tau in regulating Arc, a key regulator of synaptic plasticity (105). These findings carry implications for both tau and Arc. For tau, it suggests a new potential mechanism through which tau regulates synaptic plasticity. While the role of Arc in regulating LTP has been brought into question (106), the role of Arc in regulating protein-synthesis dependent forms of LTD is well-established, and the dysregulation in Arc could potentially be an underlying mechanism for the observed disruptions in LTD in Tau KO mice and Alzheimer’s disease animal models (106-108). For Arc, our findings further our understanding of Arc turnover, and establishes tau as a new regulator of Arc. The inability of P301L-tau to modulate Arc highlights the importance of distinctions in downstream signaling mechanisms activated by different tau mutants involved in neurodegenerative disease.

## Experimental Procedures

### Animals

All animal care and use were carried out in accordance with the National Institutes of Health Guidelines for the Use of Animals using approved protocols by the Georgia State University Institutional Animal Care and Use Committee. Tau KO mice and control WT C57BL/6J were obtained from The Jackson Laboratories (stocks #007251 and #000664) (109). The following primers were used to validate the genotype of Tau KO and C57BL/6J mice: mutant forward: 5’-GCCAGAGGCCACTTGTGTAG-3’, WT forward: 5’-AATGGAAGACCATGCTGGAG-3’ and Common: 5’-ATTCAACCCCCTCGAATTTT-3’ according to the protocol recommended by the Jackson Laboratory, with the Tau KO band at ∼170 bp, Heterozygote ∼170 bp and 269 bp and Wild type at 269 bp. Animals used in fractionation experiments and primary hippocampal cultures are balanced for sex and littermates from heterozygous pairings.

### HEK293 cell line cultures and transfections

HEK293 cells were generously provided by Dr. Jun Yin (Georgia State University). Cells were maintained in Dulbecco’s Modified Eagle Medium (DMEM; Corning # 10013CV) with 10% FBS and 1% penicillin-streptomycin (ThermoFisher). Cells were transfected at 60-70% confluency with Lipofectamine 3000 (ThermoFisher) according to the manufacturer instructions. To control for transfection efficiency, cells were plated at the same density between different conditions and the plasmid pcDNA3.1 was added with single-transfections to counterbalance other conditions that had multiple DNAs that were transfected. Culture media was exchanged 4 hr post-transfection to remove transfection material and cells were harvested 48 h later. For proteasome inhibition experiments, cells were treated with 10 μM MG-132 (sigma # 474790) for 4 hr before harvesting. MG-132 is an aldehyde peptide and a potent proteasome inhibitor that blocks the proteasome by forming a hemiacetal with the hydroxyl of the 20S active site threonines (80). For lysosome inhibition experiments, cells were treated with 50 μM leupeptin (Sigma # L2884) and 10 mM NH_4_Cl (Sigma # A9434) as described in (41) for 6 hr before harvesting. For GSK3α/β inhibition, cultures were treated with 1-2 μM CHIR 98014 (Tocris #6695) as described in (44) for 4 hr before harvesting. Plasmids used for transfection include the following: pcDNA 3.1, pEGFP-C3 (Clontech), pCMV-tdTomato, pRK5-myc-Arc (generously provided by Dr. Paul Worley, Johns Hopkins University), pRK5-myc-Arc5KR, pRK5-myc-Arc S170A/T175A, pRK5-myc-Arc T368A, pRK5-myc-Arc T380A, pRK5-myc-Arc K92Q, pRK5-myc-Arc Δ C-terminal, pRK5-myc-Arc Δ CC, pRK5-myc-Arc Δ EB (36), pEGFP-WT Tau (Addgene #46904) and pEGFP-P301L tau (Addgene #46908).

### Cloning

QuickChange™ Site-directed mutagenesis (SDM) was used to generate the mutants pRK5-myc-Arc S170A/T175A, pRK5-myc-Arc T368A, pRK5-myc-Arc T380A, pRK5-myc-Arc K92Q from the pRK5-myc-Arc backbone. Primers used were as follows: S170A/T175A For.: 5’-GGCTACGACTACACTGTTGCCCCCTATGCCATCGCCCCGCCACCTGCCGCAGGA-3’ S170A/T175A Rev.: 5’-TCCTGCGGCAGGTGGCGGGGCGATGGCATAGGGGGCAACAGTGTAGTCGTAGC-3’ T368A For.: 5’-GGCAGCTGAGCCTTCTGTCGCCCCTCTGCCCACAGAGGATG-3’ T368A Rev.: 5’-CATCCTCTGTGGGCAGAGGGGCGACAGAAGGCTCAGCTGCC-3’ K92Q For.: 5’-GGAAGAAGTCCATCCAGGCCTGTCTCTGC-3’ K92Q Rev.: 5’-GCAGAGACAGGCCTGGATGGACTTCTTCC-3’ The T380A mutant was generated using the overlap extension method as previously described in (110) with the flanking primers at the site of the mutation. T380A 3’5’ flanking primer: 5’-GAAGTCGACCCCGGGAATGGAGCTGGA-3’ T380A 5’3’ flanking primer: 5’ GAAGGATCCTTACTTACTTAGCGGCCG 3’ Fwd.: 5’-GATGAGACTGGGGCACTCGCCCCTGCTCTTACCAGCGAG-3’ Rev.: 5’-CTCGCTGGTAAGAGCAGGGGCGAGTGCCCCAGTCTCATC-3’ BamHI and SalI restriction enzymes (New England Biolabs) were used to subclone the mutated fragment into the pRK5-myc-Arc backbone.

### Western Blotting

HEK293 Cells were harvested and then cell pellets were lysed on ice in radioimmunoprecipitation assay (RIPA) buffer (150 mM NaCl, 50 mM Tris-HCl, 1% v/v Nonidet P-40, 0.5% sodium deoxycholate, 0.1% SDS) with 1 mM DTT and protease inhibitors (0.1 mM PMSF, 1 μM leupeptin, 0.15 μM aprotinin). Lysates were centrifuged at 13,000 rpm for 20 min at 4°C to precipitate insoluble extracts. Protein concentrations were measured using the Pierce 660 assay (ThermoFisher). To extract insoluble proteins, the precipitated pellet was resuspended in RIPA buffer then centrifuged at 13,000 rpm for 20 min at 4°C then resuspended in 70% formic acid and vortexed for 2 min. The samples were centrifuged at 13,000 rpm for 20 min at 4°C, the pellet discarded and 20 volumes of neutralization buffer (1M Tris Base, 0.5 M Na_2_PO_4_ with protease and phosphatase inhibitors) were added to the supernatant. Proteins were separated by SDS-PAGE then transferred to nitrocellulose membrane at 4°C (0.45 μm pore size, Bio-Rad). Membranes were blocked overnight at 4° in Intercept tris-buffered saline (TBS) blocking buffer (LI-COR) then incubated in primary antibodies in 1:1 blocking buffer to 1% Tween-20 in TBS (TBST) with 0.02% NaN_3_ overnight at 4°C. Membranes were washed 3 times with double distilled water for 5 min, and secondary antibodies in 1:1 blocking buffer to TBST 0.1% SDS were added to the membranes for 1 hr at room temperature, then washed 2 times with TBST and 1 time with double distilled water for 5 min.

Primary antibodies used: mouse anti-myc (Santa Cruz #sc-40) at 1:1000, rabbit anti-GFP (Novus Biologicals # NB600-308) at 1:1000, mouse anti-β Actin (Genetex #GTX629630) at 1:3000, mouse anti-Tau-1 (Fisher #MAB3420MI) at 1:1000, mouse anti-Tubulin (Genetex # GT114) at 1:1000, mouse anti-Ubiquitin (Sigma # U5379) at 1:500, rabbit anti-LC3 (Novus biologicals #NB100-2220) at 1:500, rabbit anti-p62 (Proteintech #18420-1-AP) at 1:1000.

Secondary antibodies used: IRDye goat anti-mouse 680RD (LI-COR #926-68070) at 1:20,000; IRDye donkey anti-rabbit 680RD (LI-COR # 925-68073) at 1:20,000; IRDye donkey anti-mouse 800CW (LI-COR #926-32212) at 1:15,000; IRDye goat anti-rabbit 800CW (LI-COR # 926-32211) at 1:15,000.

Western blot membranes were scanned using the LI-COR Odyssey CLx scanner (low scan quality, 163 μm scan resolution, auto channel intensities). Images were analyzed using ImageJ software (NIH) with the Gel Analysis tool or Image studio Lite software (Li-COR Biosciences). To adjust high background, the Subtract Background tool in FIJI was used on the whole channel with a rolling ball radius of 20-50 pixels.

### Co-Immunoprecipitation

Transfected cells were lysed in IP buffer (20 mM Tris-HCl, 3 mM EDTA, 3 mM EGTA, 150 mM NaCl, 1% Triton X-100, pH 7.4) with 1 mM DTT, protease and phosphatase inhibitors (0.1 mM PMSF, 1 μM leupeptin, 0.15 μM aprotinin, and 1:2000 Halt phosphatase inhibitor cocktail; ThermoFisher #78420). Lysates were centrifuged at 13,000 rpm for 20 min at 4°C to precipitate insoluble extracts. Protein concentration was determined using the Pierce assay 660 assay (ThermoFisher). 1 mg of protein was used for each condition and brought up to a total volume of 1 ml in IP buffer. Beads (Protein A/G PLUS-Agarose; Santa Cruz #sc-2003) were pre-equilibrated in IP buffer with inhibitors. Protein samples were incubated with 2.5 μg/sample of the primary antibody (goat anti-myc (bethyl #A190-104A) or mouse anti-myc (Santa Cruz #SC-40)) and left to tumble for 1 hr at 4°C, then equal volume of beads suspension was added per sample and left to tumble overnight. Samples were centrifuged for 45 seconds at 13,000 rpm to pellet the beads. The supernatant was discarded, and beads were washed 3 times for 5 min with IP buffer before adding 2X SDS sample buffer (4% SDS, 20% glycerol, 0.2% bromophenol blue, 3% DTT, 0.1 M Tris-HCl, 1:1000 β-mercaptoethanol, pH 6.8) and heating to 45°C for 5 min. Proteins were then separated using SDS-PAGE as described above.

For the ubiquitination assay, the same protocol was followed except that RIPA buffer was used instead of IP buffer.

### Tissue Fractionation

Dissected hippocampus from male and female 3-month old WT and Tau KO mice of mixed sex were homogenized in 10 volumes of HEPES-buffered sucrose (0.32 M sucrose, 4mM HEPES, pH 7.4) with 1 mM DTT and protease inhibitors (0.1 mM PMSF, 1 μM leupeptin, 0.15 μM aprotinin). Tissue homogenate was spun at 800 x*g* for 15 min at 4°C to precipitate the nuclear fraction (P1). The resulting supernatant (S1) was spun at 10,000 x*g* for 15 min to yield the crude synaptosomal pellet (P2). P2 was washed by resuspending in 10 volumes of HEPES-buffered sucrose and re-spinning at 10,000 x*g* for 15 min. P2 was lysed by hypoosmotic shock in 9 volumes or ice cold water with inhibitors and then rapidly adjusting to 4 mM HEPES using 1 M HEPES, pH 7.4, then left to tumble at 4°C for 30 min. Samples were then centrifuged at 25,000 x*g* for 20 min to yield the supernatant S3 (synaptosomal vesicle fraction) and the pellet P3 (synaptosomal membrane fraction). P3 was resuspended in HEPES-buffered sucrose. Quantification of protein concentrations was done using the Pierce assay and 7 μg of protein/ fraction was used in western blot analysis.

### Primary Hippocampal Neuronal Cultures

Primary hippocampal neurons of mixed sex were isolated from P0-1 mice as previously described (40) and cultured on poly-d-lysine-coated coverslips (0.1 mg/ml) in 24-well plates at a density of 75,000 cells/well. Cultures were maintained in neuronal feeding media: Neurobasal media (ThermoFisher) containing 1% GlutaMAX (ThermoFisher), 2% B-27 (ThermoFisher), 4.8 μg/mL 5-Fluoro-2’-deoxyuridine (Sigma), and 0.2 μg/mL Gentamicin (Sigma). On day *in vitro* (DIV) 6, half the media was replaced with prewarmed fresh neuronal feeding media. Cultures were transfected with Lipofectamine 2000 (ThermoFisher) on DIV 9-12 as described in (36) with equal amount of cDNA transfected into all conditions. tdTomato was used as a cell fill to identify neuron morphology as described in (100,111) For proteasome inhibition experiments, cultures were treated with 0.5 μM tetrodotoxin citrate (TTX; Tocris #1069) and 10 μM MG132 for 4 hr before they were fixed.

### Immunocytochemistry

48 h after transfection, cultures were treated with 2 μM TTX for 4 h then fixed for 20 min at 4°C with 4% Sucrose/4% paraformaldehyde. Neurons were permeabilized with 0.2 % Saponin for 15 min then blocked in 10 % normal horse serum (NHS) in PBS for 1 hr at 37°C. Permeabilization was skipped for surface GluA1 labelling. Neurons were then incubated overnight in primary antibody in 3% NHS, then washed and incubated in secondary antibody at 1:1,000 and DAPI at 1:2,000 in 3% NHS for 1 h in the dark at room temperature. Coverslips were washed with phosphate buffer saline (PBS) then mounted onto slides with Fluorogel (GeneTex).

Primary Antibodies used: rabbit anti-Arc (synaptic systems #156003) at 1:500, mouse anti-GluA1 (Millipore MAB2263) at 1:150.

Secondary Antibodies used: Donkey anti-rabbit AlexaFluor 647 (ThermoFisher #A31573), Goat anti-mouse AlexaFluor 647 (ThermoFisher #A21240)

### Image Acquisition and Analysis

For primary hippocampal neurons, coverslips were imaged on a Zeiss LSM 700 confocal microscope under 40X (Arc experiments NA 1.4, Zeiss #420762-9900) or 63X (NA 1.4, Zeiss #420782-9900-000, GluA1 experiments) immersion lens. 12-step raw z-stack images were acquired with step size 0.42 μm. Acquisition parameters were kept constant between different conditions within the same experiment and samples were interleaved during imaging. Images were analyzed using ImageJ software (NIH). Regions of interest were manually outlined guided by the neuron morphology visualized by the tdTomato cell fill, and integrated density values were quantified for Arc and GluA1 in the initial 20 μm of apical dendrites. Dendritic spines were quantified manually on the GluA1-labelled neurons imaged at 63X on the z-stacks guided by neuron morphology visualized by tdTomato cell fill in ImageJ. Protrusions from a maximum of 100 μm length of apical dendrites and their main branch less than or equal to 3 μm and with an expanded head were counted as spines and the number of spines per dendrite was normalized to the length of the dendrite.

### Experimental design and statistical analysis

All experiments follow a between-subjects design. Statistical analysis was conducted using GraphPad prism as described in the text for each experiment. Non-parametric tests are used when the criteria for using parametric tests are not met. Data is represented as mean ± SEM with statistical significance set at 95%.

## Data Availability

All data and materials are available upon request from the lead contact author.

## Acknowledgments

We would like to thank the Dr. Jun Yin (Georgia state University), Dr. Todd Cohen (University of North Carolina-Chapel Hill) and Dr. Paul Worley (John Hopkins University) for providing reagents, and Mabb lab members for feedback on experiments. We would also like to thank the Neuroscience Institute confocal core staff and GSU Division of Animal Resources.

## Author Contributions

DWY designed the study, performed experiments, and wrote the manuscript. AS confirmed lysosomal inhibition, VT contributed to data analysis and ZDA prepared primary neuronal cultures. TYN contributed to experiment design. AMM supervised and designed the study and contributed to manuscript writing. All authors participated in manuscript writing and/or editing.

## Funding and Additional Information

This study was funded by the Brain & Behavior Research Foundation NARSAD Young Investigator Award (Grant 28549), NIH/NINDS grant 1R21NS116760 and NSF CAREER Award 2047700 to AMM, and NIH/NIGMS grant GM119571 to TYN. DWY was funded by a Georgia State University Second Century Initiative Neurogenomics Fellowship, Georgia State University Molecular Basis of Disease Fellowship, and Kenneth W. and Georganne F. Honeycutt Fellowship.

## Conflict of Interest

The authors declare no competing interests.

## Figure Legends

**Supplemental figure 1: Genotype validation for Tau KO mice and validation of subcellular fractionation from hippocampus**

***A,*** Example of successful genotyping of Tau KO mice.

***B,*** Representative western blot showing the absence of tau protein in Tau KO mice. Unpaired t-test t = 12.22, df = 12, p < 0.0001

***C,*** *Left,* Representative western blots showing GluA1, PSD-95 and Actin in total hippocampal lysates and subcellular fractions. Quantification of PSD-95 and GluA1 across all probed subcellular fractions. *Right,* Values are normalized to actin levels within the same fraction. One-way ANOVA for PSD-95, F (4, 29) = 1.988, p = 0.1228. Kruskall-Wallis for GluA1 p = 0.002. n = 6 mice

**Supplemental Figure 2: Overexpression of GFP-tau and GFP-P301L tau do not alter dendritic spine densities.**

Quantification of dendritic spines on apical dendrites per 1 μm. One-way ANOVA, p = 0.857. Scale bar = 5 μm

**Supplemental Figure 3: P301L-Tau overexpression does not decrease myc-Arc**

Representative western blots showing myc-Arc with increasing concentrations of GFP-P301L tau in HEK293 cells (0, 0.25, 0.5, 0.75, 1 and 1.5 μg). Actin was used as a loading control. pcDNA3.1 was used as a DNA filler to keep the amount of transfected DNA between titration conditions identical. *Bottom*, Quantification of myc-Arc normalized to actin showing no significant differences in Arc with increasing P301L Tau. One-way ANOVA, F (5,18) = 0.3018, p = 0.3. n= 4.

**Supplemental Figure 4: Tau does not promote lysosomal degradation of Arc**

***A,*** Representative western blots showing myc-Arc expressed alone or with GFP-tau. Cells were treated with Vehicle (water) or a combination of leupeptin (50 μM) and ammonium (10 mM) chloride for 6 hr to block lysosome degradation. Actin was used as a loading control. Blots showing markers for lysosome inhibition, LC3 and p62/SQSTM1 with total protein stain used as a loading control.

***B,*** Quantification of LC3 and p62/SQSTM1 showing a significant increase of LC3 (unpaired t-test, t = 12.52, df = 22, ****p < 0.0001) and p62/SQSTM1 (unpaired t-test, t = 10.54, df = 22, ****p < 0.0001) in cells from all conditions treated with inhibitors. n = 12.

***C,*** Quantification of myc-Arc showing a significant decrease with co-expression of GFP-tau in Vehicle and following lysosome inhibition. t-test for vehicle control, t = 10.6, df = 10, ****p < 0.0001; unpaired t-test for inhibitors, t = 5.585, df = 10, p = 0.0002. n = 6.

***D,*** Representative western blots showing RIPA-soluble and insoluble fractions of myc-Arc expressed alone or with GFP-tau.

## Notes

### Competing Interest Statement

The authors have declared no competing interest.

### Summary of Updates

We now provide new data that shows that the regulation of Arc by tau is physiological, as Tau knockout mice exhibit elevations in Arc in total hippocampal extracts, crude hippocampal synaptosomal fractions and also in dendrites of Tau knockout hippocampal neurons. We further provide details and new experiments on essential controls related to Tau-dependent Arc stability in nonneuronal cells.

## References

1. Williams, D. R. (2006) Tauopathies: classification and clinical update on neurodegenerative diseases associated with microtubule-associated protein tau. Intern Med J 36, 652–660

2. Kosik, K. S., Joachim, C. L., and Selkoe, D. J. (1986) Microtubule-associated protein tau (tau) is a major antigenic component of paired helical filaments in Alzheimer disease. Proc Natl Acad Sci U S A 83, 4044–4048

3. Wood, J. G., Mirra, S. S., Pollock, N. J., and Binder, L. I. (1986) Neurofibrillary tangles of Alzheimer disease share antigenic determinants with the axonal microtubule-associated protein tau (tau). Proc Natl Acad Sci U S A 83, 4040–4043

4. Braak, H., and Braak, E. (1995) Staging of Alzheimer’s disease-related neurofibrillary changes. Neurobiol Aging 16, 271–278; discussion 278-284

5. Andreadis, A., Brown, W. M., and Kosik, K. S. (1992) Structure and novel exons of the human tau gene. Biochemistry 31, 10626–10633

6. Wolfe, M. S. (2009) Tau mutations in neurodegenerative diseases. J Biol Chem 284, 6021–6025

7. Goedert, M., Wischik, C. M., Crowther, R. A., Walker, J. E., and Klug, A. (1988) Cloning and sequencing of the cDNA encoding a core protein of the paired helical filament of Alzheimer disease: identification as the microtubule-associated protein tau. Proc Natl Acad Sci U S A 85, 4051–4055

8. Goedert, M., Spillantini, M. G., Jakes, R., Rutherford, D., and Crowther, R. A. (1989) Multiple isoforms of human microtubule-associated protein tau: sequences and localization in neurofibrillary tangles of Alzheimer’s disease. Neuron 3, 519–526

9. Reynolds, C. H., Betts, J. C., Blackstock, W. P., Nebreda, A. R., and Anderton, B. H. (2000) Phosphorylation sites on tau identified by nanoelectrospray mass spectrometry: differences in vitro between the mitogen-activated protein kinases ERK2, c-Jun N-terminal kinase and P38, and glycogen synthase kinase-3beta. J Neurochem 74, 1587–1595

10. Min, S. W., Cho, S. H., Zhou, Y., Schroeder, S., Haroutunian, V., Seeley, W. W., Huang, E. J., Shen, Y., Masliah, E., Mukherjee, C., Meyers, D., Cole, P. A., Ott, M., and Gan, L. (2010) Acetylation of tau inhibits its degradation and contributes to tauopathy. Neuron 67, 953–966

11. Cohen, T. J., Guo, J. L., Hurtado, D. E., Kwong, L. K., Mills, I. P., Trojanowski, J. Q., and Lee, V. M. (2011) The acetylation of tau inhibits its function and promotes pathological tau aggregation. Nat Commun 2, 252

12. Shimura, H., Schwartz, D., Gygi, S. P., and Kosik, K. S. (2004) CHIP-Hsc70 complex ubiquitinates phosphorylated tau and enhances cell survival. J Biol Chem 279, 4869–4876

13. Tuerde, D., Kimura, T., Miyasaka, T., Furusawa, K., Shimozawa, A., Hasegawa, M., Ando, K., and Hisanaga, S. I. (2018) Isoform-independent and -dependent phosphorylation of microtubule-associated protein tau in mouse brain during postnatal development. J Biol Chem 293, 1781–1793

14. Wang, Y., and Mandelkow, E. (2016) Tau in physiology and pathology. Nat Rev Neurosci 17, 5–21

15. Harrison, T. M., La Joie, R., Maass, A., Baker, S. L., Swinnerton, K., Fenton, L., Mellinger, T. J., Edwards, L., Pham, J., Miller, B. L., Rabinovici, G. D., and Jagust, W. J. (2019) Longitudinal tau accumulation and atrophy in aging and alzheimer disease. Ann Neurol 85, 229–240

16. Torres, A. K., Rivera, B. I., Polanco, C. M., Jara, C., and Tapia-Rojas, C. (2022) Phosphorylated tau as a toxic agent in synaptic mitochondria: implications in aging and Alzheimer’s disease. Neural Regen Res 17, 1645–1651

17. Strang, K. H., Golde, T. E., and Giasson, B. I. (2019) MAPT mutations, tauopathy, and mechanisms of neurodegeneration. Lab Invest 99, 912–928

18. D’Souza, I., and Schellenberg, G. D. (2005) Regulation of tau isoform expression and dementia. Biochim Biophys Acta 1739, 104–115

19. Hutton, M., Lendon, C. L., Rizzu, P., Baker, M., Froelich, S., Houlden, H., Pickering-Brown, S., Chakraverty, S., Isaacs, A., Grover, A., Hackett, J., Adamson, J., Lincoln, S., Dickson, D., Davies, P., Petersen, R. C., Stevens, M., de Graaff, E., Wauters, E., van Baren, J., Hillebrand, M., Joosse, M., Kwon, J. M., Nowotny, P., Che, L. K., Norton, J., Morris, J. C., Reed, L. A., Trojanowski, J., Basun, H., Lannfelt, L., Neystat, M., Fahn, S., Dark, F., Tannenberg, T., Dodd, P. R., Hayward, N., Kwok, J. B., Schofield, P. R., Andreadis, A., Snowden, J., Craufurd, D., Neary, D., Owen, F., Oostra, B. A., Hardy, J., Goate, A., van Swieten, J., Mann, D., Lynch, T., and Heutink, P. (1998) Association of missense and 5’-splice-site mutations in tau with the inherited dementia FTDP-17. Nature 393, 702–705

20. Barbier, P., Zejneli, O., Martinho, M., Lasorsa, A., Belle, V., Smet-Nocca, C., Tsvetkov, P. O., Devred, F., and Landrieu, I. (2019) Role of Tau as a Microtubule-Associated Protein: Structural and Functional Aspects. Front Aging Neurosci 11, 204

21. Sotiropoulos, I., Galas, M. C., Silva, J. M., Skoulakis, E., Wegmann, S., Maina, M. B., Blum, D., Sayas, C. L., Mandelkow, E. M., Mandelkow, E., Spillantini, M. G., Sousa, N., Avila, J., Medina, M., Mudher, A., and Buee, L. (2017) Atypical, non-standard functions of the microtubule associated Tau protein. Acta Neuropathol Commun 5, 91

22. Liu, C., and Gotz, J. (2013) Profiling murine tau with 0N, 1N and 2N isoform-specific antibodies in brain and peripheral organs reveals distinct subcellular localization, with the 1N isoform being enriched in the nucleus. PLoS One 8, e84849

23. Kimura, T., Whitcomb, D. J., Jo, J., Regan, P., Piers, T., Heo, S., Brown, C., Hashikawa, T., Murayama, M., Seok, H., Sotiropoulos, I., Kim, E., Collingridge, G. L., Takashima, A., and Cho, K. (2014) Microtubule-associated protein tau is essential for long-term depression in the hippocampus. Philos Trans R Soc Lond B Biol Sci 369, 20130144

24. Ahmed, T., Van der Jeugd, A., Blum, D., Galas, M. C., D’Hooge, R., Buee, L., and Balschun, D. (2014) Cognition and hippocampal synaptic plasticity in mice with a homozygous tau deletion. Neurobiol Aging 35, 2474–2478

25. Prikas, E., Paric, E., Asih, P. R., Stefanoska, K., Stefen, H., Fath, T., Poljak, A., and Ittner, A. (2022) Tau target identification reveals NSF-dependent effects on AMPA receptor trafficking and memory formation. EMBO J 41, e10242

26. Trepanier, C. H., Jackson, M. F., and MacDonald, J. F. (2012) Regulation of NMDA receptors by the tyrosine kinase Fyn. FEBS J 279, 12–19

27. Waung, M. W., Pfeiffer, B. E., Nosyreva, E. D., Ronesi, J. A., and Huber, K. M. (2008) Rapid translation of Arc/Arg3.1 selectively mediates mGluR-dependent LTD through persistent increases in AMPAR endocytosis rate. Neuron 59, 84–97

28. Rial Verde, E. M., Lee-Osbourne, J., Worley, P. F., Malinow, R., and Cline, H. T. (2006) Increased expression of the immediate-early gene arc/arg3.1 reduces AMPA receptor-mediated synaptic transmission. Neuron 52, 461–474

29. Messaoudi, E., Kanhema, T., Soule, J., Tiron, A., Dagyte, G., da Silva, B., and Bramham, C. R. (2007) Sustained Arc/Arg3.1 synthesis controls long-term potentiation consolidation through regulation of local actin polymerization in the dentate gyrus in vivo. J Neurosci 27, 10445–10455

30. Shepherd, J. D., Rumbaugh, G., Wu, J., Chowdhury, S., Plath, N., Kuhl, D., Huganir, R. L., and Worley, P. F. (2006) Arc/Arg3.1 mediates homeostatic synaptic scaling of AMPA receptors. Neuron 52, 475–484

31. Okuno, H., Akashi, K., Ishii, Y., Yagishita-Kyo, N., Suzuki, K., Nonaka, M., Kawashima, T., Fujii, H., Takemoto-Kimura, S., Abe, M., Natsume, R., Chowdhury, S., Sakimura, K., Worley, P. F., and Bito, H. (2012) Inverse synaptic tagging of inactive synapses via dynamic interaction of Arc/Arg3.1 with CaMKIIbeta. Cell 149, 886–898

32. Chowdhury, S., Shepherd, J. D., Okuno, H., Lyford, G., Petralia, R. S., Plath, N., Kuhl, D., Huganir, R. L., and Worley, P. F. (2006) Arc/Arg3.1 interacts with the endocytic machinery to regulate AMPA receptor trafficking. Neuron 52, 445–459

33. DaSilva, L. L., Wall, M. J., L, P. d. A., Wauters, S. C., Januario, Y. C., Muller, J., and Correa, S. A. (2016) Activity-Regulated Cytoskeleton-Associated Protein Controls AMPAR Endocytosis through a Direct Interaction with Clathrin-Adaptor Protein 2. eNeuro 3

34. Guzowski, J. F., McNaughton, B. L., Barnes, C. A., and Worley, P. F. (1999) Environment-specific expression of the immediate-early gene Arc in hippocampal neuronal ensembles. Nat Neurosci 2, 1120–1124

35. Greer, P. L., Hanayama, R., Bloodgood, B. L., Mardinly, A. R., Lipton, D. M., Flavell, S. W., Kim, T. K., Griffith, E. C., Waldon, Z., Maehr, R., Ploegh, H. L., Chowdhury, S., Worley, P. F., Steen, J., and Greenberg, M. E. (2010) The Angelman Syndrome protein Ube3A regulates synapse development by ubiquitinating arc. Cell 140, 704–716

36. Mabb, A. M., Je, H. S., Wall, M. J., Robinson, C. G., Larsen, R. S., Qiang, Y., Correa, S. A., and Ehlers, M. D. (2014) Triad3A regulates synaptic strength by ubiquitination of Arc. Neuron 82, 1299–1316

37. Mabb, A. M., and Ehlers, M. D. (2018) Arc ubiquitination in synaptic plasticity. Semin Cell Dev Biol 77, 10–16

38. Liu, Y., Sun, Y., Huang, Y., Cheng, K., Xu, Y., Tian, Q., and Zhang, S. (2021) CHIP promotes Wnt signaling and regulates Arc stability by recruiting and polyubiquitinating LEF1 or Arc. Cell Death Discov 7, 5

39. Rao, V. R., Pintchovski, S. A., Chin, J., Peebles, C. L., Mitra, S., and Finkbeiner, S. (2006) AMPA receptors regulate transcription of the plasticity-related immediate-early gene Arc. Nat Neurosci 9, 887–895

40. Wall, M. J., Collins, D. R., Chery, S. L., Allen, Z. D., Pastuzyn, E. D., George, A. J., Nikolova, V. D., Moy, S. S., Philpot, B. D., Shepherd, J. D., Muller, J., Ehlers, M. D., Mabb, A. M., and Correa, S. A. L. (2018) The Temporal Dynamics of Arc Expression Regulate Cognitive Flexibility. Neuron 98, 1124–1132 e1127

41. Yan, J., Porch, M. W., Court-Vazquez, B., Bennett, M. V. L., and Zukin, R. S. (2018) Activation of autophagy rescues synaptic and cognitive deficits in fragile X mice. Proc Natl Acad Sci U S A 115, E9707–E9716

42. Ramachandran, K. V., Fu, J. M., Schaffer, T. B., Na, C. H., Delannoy, M., and Margolis, S. S. (2018) Activity-Dependent Degradation of the Nascentome by the Neuronal Membrane Proteasome. Mol Cell 71, 169–177 e166

43. Ramachandran, K. V., and Margolis, S. S. (2017) A mammalian nervous-system-specific plasma membrane proteasome complex that modulates neuronal function. Nat Struct Mol Biol 24, 419–430

44. Gozdz, A., Nikolaienko, O., Urbanska, M., Cymerman, I. A., Sitkiewicz, E., Blazejczyk, M., Dadlez, M., Bramham, C. R., and Jaworski, J. (2017) GSK3alpha and GSK3beta Phosphorylate Arc and Regulate its Degradation. Front Mol Neurosci 10, 192

45. Nair, R. R., Patil, S., Tiron, A., Kanhema, T., Panja, D., Schiro, L., Parobczak, K., Wilczynski, G., and Bramham, C. R. (2017) Dynamic Arc SUMOylation and Selective Interaction with F-Actin-Binding Protein Drebrin A in LTP Consolidation In Vivo. Front Synaptic Neurosci 9, 8

46. Bramham, C. R., Alme, M. N., Bittins, M., Kuipers, S. D., Nair, R. R., Pai, B., Panja, D., Schubert, M., Soule, J., Tiron, A., and Wibrand, K. (2010) The Arc of synaptic memory. Exp Brain Res 200, 125–140

47. Craig, T. J., Jaafari, N., Petrovic, M. M., Jacobs, S. C., Rubin, P. P., Mellor, J. R., and Henley, J. M. (2012) Homeostatic synaptic scaling is regulated by protein SUMOylation. J Biol Chem 287, 22781–22788

48. Barylko, B., Wilkerson, J. R., Cavalier, S. H., Binns, D. D., James, N. G., Jameson, D. M., Huber, K. M., and Albanesi, J. P. (2018) Palmitoylation and Membrane Binding of Arc/Arg3.1: A Potential Role in Synaptic Depression. Biochemistry 57, 520–524

49. Lalonde, J., Reis, S. A., Sivakumaran, S., Holland, C. S., Wesseling, H., Sauld, J. F., Alural, B., Zhao, W. N., Steen, J. A., and Haggarty, S. J. (2017) Chemogenomic analysis reveals key role for lysine acetylation in regulating Arc stability. Nat Commun 8, 1659

50. Wegenast-Braun, B. M., Fulgencio Maisch, A., Eicke, D., Radde, R., Herzig, M. C., Staufenbiel, M., Jucker, M., and Calhoun, M. E. (2009) Independent effects of intra- and extracellular Abeta on learning-related gene expression. Am J Pathol 175, 271–282

51. Dickey, C. A., Gordon, M. N., Mason, J. E., Wilson, N. J., Diamond, D. M., Guzowski, J. F., and Morgan, D. (2004) Amyloid suppresses induction of genes critical for memory consolidation in APP + PS1 transgenic mice. J Neurochem 88, 434–442

52. Dickey, C. A., Loring, J. F., Montgomery, J., Gordon, M. N., Eastman, P. S., and Morgan, D. (2003) Selectively reduced expression of synaptic plasticity-related genes in amyloid precursor protein + presenilin-1 transgenic mice. J Neurosci 23, 5219–5226

53. Perez-Cruz, C., Nolte, M. W., van Gaalen, M. M., Rustay, N. R., Termont, A., Tanghe, A., Kirchhoff, F., and Ebert, U. (2011) Reduced spine density in specific regions of CA1 pyramidal neurons in two transgenic mouse models of Alzheimer’s disease. J Neurosci 31, 3926–3934

54. Wu, J., Petralia, R. S., Kurushima, H., Patel, H., Jung, M. Y., Volk, L., Chowdhury, S., Shepherd, J. D., Dehoff, M., Li, Y., Kuhl, D., Huganir, R. L., Price, D. L., Scannevin, R., Troncoso, J. C., Wong, P. C., and Worley, P. F. (2011) Arc/Arg3.1 regulates an endosomal pathway essential for activity-dependent beta-amyloid generation. Cell 147, 615–628

55. Ittner, L. M., Ke, Y. D., Delerue, F., Bi, M., Gladbach, A., van Eersel, J., Wolfing, H., Chieng, B. C., Christie, M. J., Napier, I. A., Eckert, A., Staufenbiel, M., Hardeman, E., and Gotz, J. (2010) Dendritic function of tau mediates amyloid-beta toxicity in Alzheimer’s disease mouse models. Cell 142, 387–397

56. Holth, J. K., Bomben, V. C., Reed, J. G., Inoue, T., Younkin, L., Younkin, S. G., Pautler, R. G., Botas, J., and Noebels, J. L. (2013) Tau loss attenuates neuronal network hyperexcitability in mouse and Drosophila genetic models of epilepsy. J Neurosci 33, 1651–1659

57. Roberson, E. D., Scearce-Levie, K., Palop, J. J., Yan, F., Cheng, I. H., Wu, T., Gerstein, H., Yu, G. Q., and Mucke, L. (2007) Reducing endogenous tau ameliorates amyloid beta-induced deficits in an Alzheimer’s disease mouse model. Science 316, 750–754

58. Peebles, C. L., Yoo, J., Thwin, M. T., Palop, J. J., Noebels, J. L., and Finkbeiner, S. (2010) Arc regulates spine morphology and maintains network stability in vivo. Proc Natl Acad Sci U S A 107, 18173–18178

59. van Kerkhof, P., Alves dos Santos, C. M., Sachse, M., Klumperman, J., Bu, G., and Strous, G. J. (2001) Proteasome inhibitors block a late step in lysosomal transport of selected membrane but not soluble proteins. Mol Biol Cell 12, 2556–2566

60. Shroff, A., and Nazarko, T. Y. (2022) SQSTM1, lipid droplets and current state of their lipophagy affairs. Autophagy, 1-4

61. Lippens, G., Sillen, A., Landrieu, I., Amniai, L., Sibille, N., Barbier, P., Leroy, A., Hanoulle, X., and Wieruszeski, J. M. (2007) Tau aggregation in Alzheimer’s disease: what role for phosphorylation? Prion 1, 21–25

62. Ackmann, M., Wiech, H., and Mandelkow, E. (2000) Nonsaturable binding indicates clustering of tau on the microtubule surface in a paired helical filament-like conformation. J Biol Chem 275, 30335–30343

63. Ishihara, T., Higuchi, M., Zhang, B., Yoshiyama, Y., Hong, M., Trojanowski, J. Q., and Lee, V. M. (2001) Attenuated neurodegenerative disease phenotype in tau transgenic mouse lacking neurofilaments. J Neurosci 21, 6026–6035

64. Dou, F., Netzer, W. J., Tanemura, K., Li, F., Hartl, F. U., Takashima, A., Gouras, G. K., Greengard, P., and Xu, H. (2003) Chaperones increase association of tau protein with microtubules. Proc Natl Acad Sci U S A 100, 721–726

65. Palop, J. J., Chin, J., Bien-Ly, N., Massaro, C., Yeung, B. Z., Yu, G. Q., and Mucke, L. (2005) Vulnerability of dentate granule cells to disruption of arc expression in human amyloid precursor protein transgenic mice. J Neurosci 25, 9686–9693

66. Parra-Damas, A., Valero, J., Chen, M., Espana, J., Martin, E., Ferrer, I., Rodriguez-Alvarez, J., and Saura, C. A. (2014) Crtc1 activates a transcriptional program deregulated at early Alzheimer’s disease-related stages. J Neurosci 34, 5776–5787

67. Xu, C., Huang, H., Zhang, M., Zhang, P., Li, Z., Liu, X., and Fang, M. (2022) Methyltransferase-Like 3 Rescues the Amyloid-beta protein-Induced Reduction of Activity-Regulated Cytoskeleton Associated Protein Expression via YTHDF1-Dependent N6-Methyladenosine Modification. Front Aging Neurosci 14, 890134

68. Privitera, L., Hogg, E. L., Lopes, M., Domingos, L. B., Gaestel, M., Muller, J., Wall, M. J., and Correa, S. A. L. (2022) The MK2 cascade mediates transient alteration in mGluR-LTD and spatial learning in a murine model of Alzheimer’s disease. Aging Cell 21, e13717

69. Naert, G., and Rivest, S. (2012) Age-related changes in synaptic markers and monocyte subsets link the cognitive decline of APP(Swe)/PS1 mice. Front Cell Neurosci 6, 51

70. Rudinskiy, N., Hawkes, J. M., Betensky, R. A., Eguchi, M., Yamaguchi, S., Spires-Jones, T. L., and Hyman, B. T. (2012) Orchestrated experience-driven Arc responses are disrupted in a mouse model of Alzheimer’s disease. Nat Neurosci 15, 1422–1429

71. Rudinskiy, N., Hawkes, J. M., Wegmann, S., Kuchibhotla, K. V., Muzikansky, A., Betensky, R. A., Spires-Jones, T. L., and Hyman, B. T. (2014) Tau pathology does not affect experience-driven single-neuron and network-wide Arc/Arg3.1 responses. Acta Neuropathol Commun 2, 63

72. Schulz, L., Ramirez, P., Lemieux, A., Gonzalez, E., Thomson, T., and Frost, B. (2022) Tau-Induced Elevation of the Activity-Regulated Cytoskeleton Associated Protein Arc1 Causally Mediates Neurodegeneration in the Adult Drosophila Brain. Neuroscience

73. Steward, O., Wallace, C. S., Lyford, G. L., and Worley, P. F. (1998) Synaptic activation causes the mRNA for the IEG Arc to localize selectively near activated postsynaptic sites on dendrites. Neuron 21, 741–751

74. Moga, D. E., Calhoun, M. E., Chowdhury, A., Worley, P., Morrison, J. H., and Shapiro, M. L. (2004) Activity-regulated cytoskeletal-associated protein is localized to recently activated excitatory synapses. Neuroscience 125, 7–11

75. Ramirez-Amaya, V., Vazdarjanova, A., Mikhael, D., Rosi, S., Worley, P. F., and Barnes, C. A. (2005) Spatial exploration-induced Arc mRNA and protein expression: evidence for selective, network-specific reactivation. J Neurosci 25, 1761–1768

76. Petrucelli, L., Dickson, D., Kehoe, K., Taylor, J., Snyder, H., Grover, A., De Lucia, M., McGowan, E., Lewis, J., Prihar, G., Kim, J., Dillmann, W. H., Browne, S. E., Hall, A., Voellmy, R., Tsuboi, Y., Dawson, T. M., Wolozin, B., Hardy, J., and Hutton, M. (2004) CHIP and Hsp70 regulate tau ubiquitination, degradation and aggregation. Hum Mol Genet 13, 703–714

77. Eriksen, M. S., Nikolaienko, O., Hallin, E. I., Grodem, S., Bustad, H. J., Flydal, M. I., Merski, I., Hosokawa, T., Lascu, D., Akerkar, S., Cuellar, J., Chambers, J. J., O’Connell, R., Muruganandam, G., Loris, R., Touma, C., Kanhema, T., Hayashi, Y., Stratton, M. M., Valpuesta, J. M., Kursula, P., Martinez, A., and Bramham, C. R. (2021) Arc self-association and formation of virus-like capsids are mediated by an N-terminal helical coil motif. FEBS J 288, 2930–2955

78. Ashley, J., Cordy, B., Lucia, D., Fradkin, L. G., Budnik, V., and Thomson, T. (2018) Retrovirus-like Gag Protein Arc1 Binds RNA and Traffics across Synaptic Boutons. Cell 172, 262–274 e211

79. Pastuzyn, E. D., Day, C. E., Kearns, R. B., Kyrke-Smith, M., Taibi, A. V., McCormick, J., Yoder, N., Belnap, D. M., Erlendsson, S., Morado, D. R., Briggs, J. A. G., Feschotte, C., and Shepherd, J. D. (2018) The Neuronal Gene Arc Encodes a Repurposed Retrotransposon Gag Protein that Mediates Intercellular RNA Transfer. Cell 173, 275

80. Kisselev, A. F., van der Linden, W. A., and Overkleeft, H. S. (2012) Proteasome inhibitors: an expanding army attacking a unique target. Chem Biol 19, 99–115

81. Morozov, A. V., and Karpov, V. L. (2019) Proteasomes and Several Aspects of Their Heterogeneity Relevant to Cancer. Front Oncol 9, 761

82. Turker, F., Cook, E. K., and Margolis, S. S. (2021) The proteasome and its role in the nervous system. Cell Chem Biol 28, 903–917

83. Boratyn, G. M., Schaffer, A. A., Agarwala, R., Altschul, S. F., Lipman, D. J., and Madden, T. L. (2012) Domain enhanced lookup time accelerated BLAST. Biol Direct 7, 12

84. Hallin, E. I., Eriksen, M. S., Baryshnikov, S., Nikolaienko, O., Grodem, S., Hosokawa, T., Hayashi, Y., Bramham, C. R., and Kursula, P. (2018) Structure of monomeric full-length ARC sheds light on molecular flexibility, protein interactions, and functional modalities. J Neurochem 147, 323–343

85. Liu, C., Song, X., Nisbet, R., and Gotz, J. (2016) Co-immunoprecipitation with Tau Isoform-specific Antibodies Reveals Distinct Protein Interactions and Highlights a Putative Role for 2N Tau in Disease. J Biol Chem 291, 8173–8188

86. Hori, T., Eguchi, K., Wang, H. Y., Miyasaka, T., Guillaud, L., Taoufiq, Z., Mahapatra, S., Yamada, H., Takei, K., and Takahashi, T. (2022) Microtubule assembly by tau impairs endocytosis and neurotransmission via dynamin sequestration in Alzheimer’s disease synapse model. Elife 11

87. Ittner, A., and Ittner, L. M. (2018) Dendritic Tau in Alzheimer’s Disease. Neuron 99, 13–27

88. Regan, P., Piers, T., Yi, J. H., Kim, D. H., Huh, S., Park, S. J., Ryu, J. H., Whitcomb, D. J., and Cho, K. (2015) Tau phosphorylation at serine 396 residue is required for hippocampal LTD. J Neurosci 35, 4804–4812

89. Suzuki, M., and Kimura, T. (2017) Microtubule-associated tau contributes to intra-dendritic trafficking of AMPA receptors in multiple ways. Neurosci Lett 653, 276–282

90. Franchini, L., Stanic, J., Ponzoni, L., Mellone, M., Carrano, N., Musardo, S., Zianni, E., Olivero, G., Marcello, E., Pittaluga, A., Sala, M., Bellone, C., Racca, C., Di Luca, M., and Gardoni, F. (2019) Linking NMDA Receptor Synaptic Retention to Synaptic Plasticity and Cognition. iScience 19, 927–939

91. Hoover, B. R., Reed, M. N., Su, J., Penrod, R. D., Kotilinek, L. A., Grant, M. K., Pitstick, R., Carlson, G. A., Lanier, L. M., Yuan, L. L., Ashe, K. H., and Liao, D. (2010) Tau mislocalization to dendritic spines mediates synaptic dysfunction independently of neurodegeneration. Neuron 68, 1067–1081

92. Gamache, J., Benzow, K., Forster, C., Kemper, L., Hlynialuk, C., Furrow, E., Ashe, K. H., and Koob, M. D. (2019) Factors other than hTau overexpression that contribute to tauopathy-like phenotype in rTg4510 mice. Nat Commun 10, 2479

93. Goodwin, L. O., Splinter, E., Davis, T. L., Urban, R., He, H., Braun, R. E., Chesler, E. J., Kumar, V., van Min, M., Ndukum, J., Philip, V. M., Reinholdt, L. G., Svenson, K., White, J. K., Sasner, M., Lutz, C., and Murray, S. A. (2019) Large-scale discovery of mouse transgenic integration sites reveals frequent structural variation and insertional mutagenesis. Genome Res 29, 494–505

94. Bachmann, S., Bell, M., Klimek, J., and Zempel, H. (2021) Differential Effects of the Six Human TAU Isoforms: Somatic Retention of 2N-TAU and Increased Microtubule Number Induced by 4R-TAU. Front Neurosci 15, 643115

95. Cherry, J. D., Esnault, C. D., Baucom, Z. H., Tripodis, Y., Huber, B. R., Alvarez, V. E., Stein, T. D., Dickson, D. W., and McKee, A. C. (2021) Tau isoforms are differentially expressed across the hippocampus in chronic traumatic encephalopathy and Alzheimer’s disease. Acta Neuropathol Commun 9, 86

96. Alonso, A. D., Zaidi, T., Novak, M., Barra, H. S., Grundke-Iqbal, I., and Iqbal, K. (2001) Interaction of tau isoforms with Alzheimer’s disease abnormally hyperphosphorylated tau and in vitro phosphorylation into the disease-like protein. J Biol Chem 276, 37967–37973

97. Goedert, M., and Jakes, R. (1990) Expression of separate isoforms of human tau protein: correlation with the tau pattern in brain and effects on tubulin polymerization. EMBO J 9, 4225–4230

98. Lee, V. M., Goedert, M., and Trojanowski, J. Q. (2001) Neurodegenerative tauopathies. Annu Rev Neurosci 24, 1121–1159

99. Hall, A. M., and Roberson, E. D. (2012) Mouse models of Alzheimer’s disease. Brain Res Bull 88, 3–12

100. Gamache, J. E., Kemper, L., Steuer, E., Leinonen-Wright, K., Choquette, J. M., Hlynialuk, C., Benzow, K., Vossel, K. A., Xia, W., Koob, M. D., and Ashe, K. H. (2020) Developmental Pathogenicity of 4-Repeat Human Tau Is Lost with the P301L Mutation in Genetically Matched Tau-Transgenic Mice. J Neurosci 40, 220–236

101. Yakout, D. W., Shree, N., and Mabb, A. M. (2021) Effect of pharmacological manipulations on Arc function. Curr Res Pharmacol Drug Discov 2, 100013

102. Plath, N., Ohana, O., Dammermann, B., Errington, M. L., Schmitz, D., Gross, C., Mao, X., Engelsberg, A., Mahlke, C., Welzl, H., Kobalz, U., Stawrakakis, A., Fernandez, E., Waltereit, R., Bick-Sander, A., Therstappen, E., Cooke, S. F., Blanquet, V., Wurst, W., Salmen, B., Bosl, M. R., Lipp, H. P., Grant, S. G., Bliss, T. V., Wolfer, D. P., and Kuhl, D. (2006) Arc/Arg3.1 is essential for the consolidation of synaptic plasticity and memories. Neuron 52, 437–444

103. Buschert, V., Bokde, A. L., and Hampel, H. (2010) Cognitive intervention in Alzheimer disease. Nat Rev Neurol 6, 508–517

104. Pluta, R., and Ulamek-Koziol, M. (2020) Tau Protein-Targeted Therapies in Alzheimer’s Disease: Current State and Future Perspectives. in Alzheimer’s Disease: Drug Discovery (Huang, X. ed.), Brisbane (AU). pp

105. Nikolaienko, O., Patil, S., Eriksen, M. S., and Bramham, C. R. (2018) Arc protein: a flexible hub for synaptic plasticity and cognition. Semin Cell Dev Biol 77, 33–42

106. Kyrke-Smith, M., Volk, L. J., Cooke, S. F., Bear, M. F., Huganir, R. L., and Shepherd, J. D. (2021) The Immediate Early Gene Arc Is Not Required for Hippocampal Long-Term Potentiation. J Neurosci 41, 4202–4211

107. Wilkerson, J. R., Albanesi, J. P., and Huber, K. M. (2018) Roles for Arc in metabotropic glutamate receptor-dependent LTD and synapse elimination: Implications in health and disease. Semin Cell Dev Biol 77, 51–62

108. Mango, D., Saidi, A., Cisale, G. Y., Feligioni, M., Corbo, M., and Nistico, R. (2019) Targeting Synaptic Plasticity in Experimental Models of Alzheimer’s Disease. Front Pharmacol 10, 778

109. Dawson, H. N., Ferreira, A., Eyster, M. V., Ghoshal, N., Binder, L. I., and Vitek, M. P. (2001) Inhibition of neuronal maturation in primary hippocampal neurons from tau deficient mice. J Cell Sci 114, 1179–1187

110. Ho, S. N., Hunt, H. D., Horton, R. M., Pullen, J. K., and Pease, L. R. (1989) Site-directed mutagenesis by overlap extension using the polymerase chain reaction. Gene 77, 51–59

111. On, V., Zahedi, A., Ethell, I. M., and Bhanu, B. (2017) Automated spatio-temporal analysis of dendritic spines and related protein dynamics. PLoS One 12, e0182958

