## Supplementary figures and images for "Tau regulates Arc stability in neuronal dendrites via a proteasome-sensitive but ubiquitin-independent pathway"

### Supplemental Fig. 1

#### Supplemental Figure 1

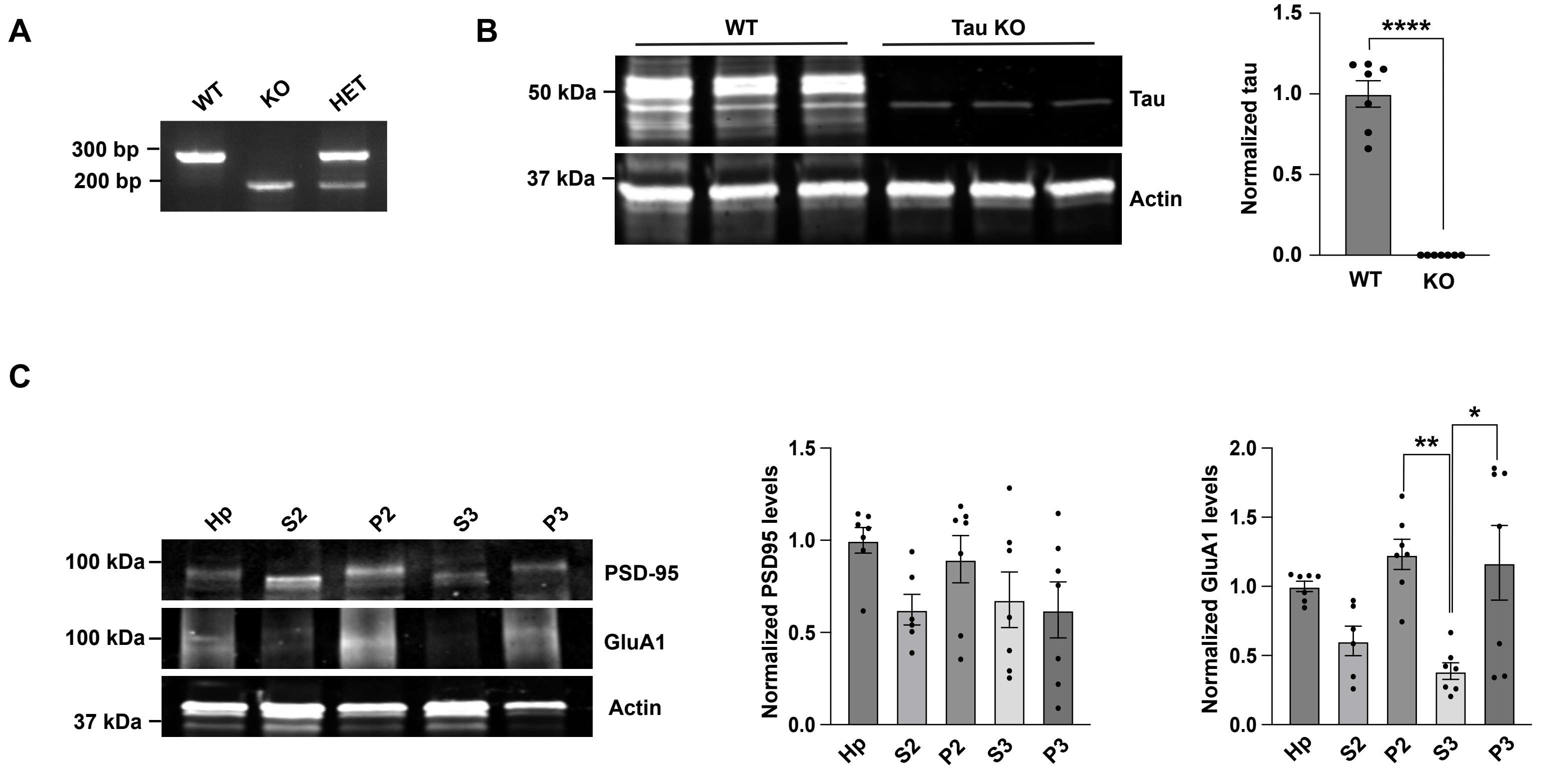

### Supplemental Fig. 2

#### Supplemental Figure 2

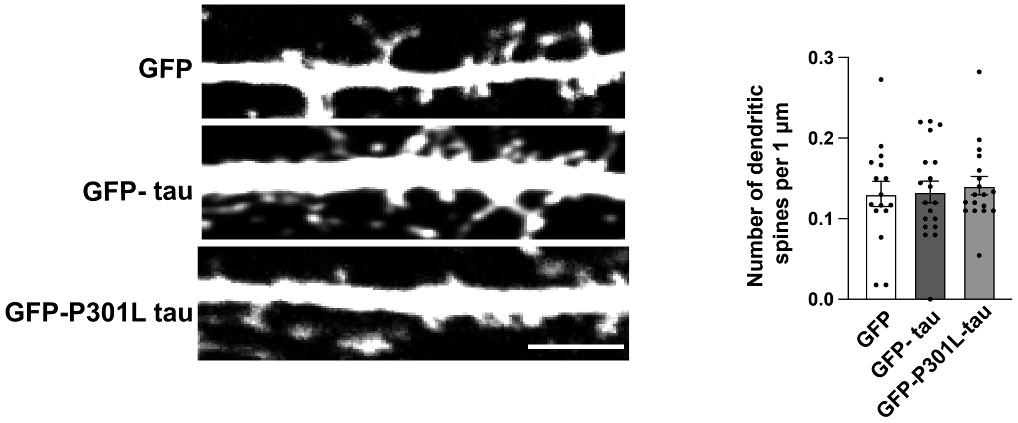

### Supplemental Fig. 3

#### Supplemental Figure 3

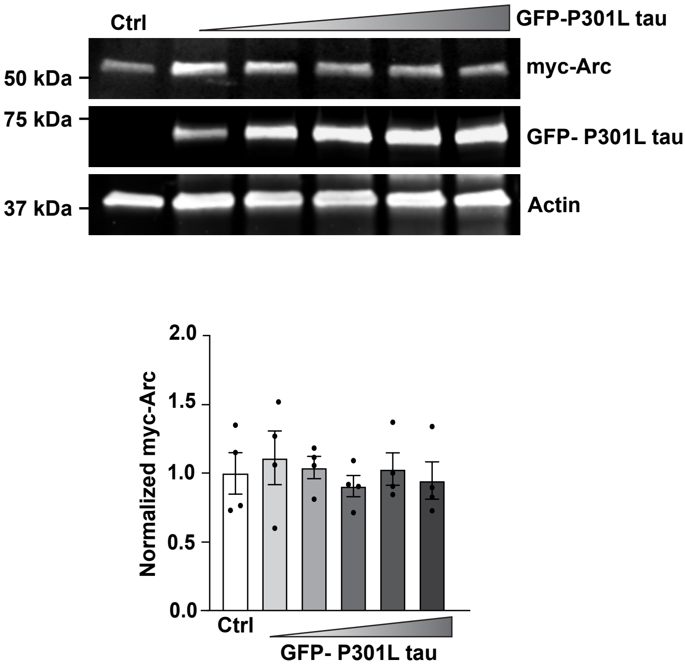

### Supplemental Fig. 4

#### Supplemental Figure 4

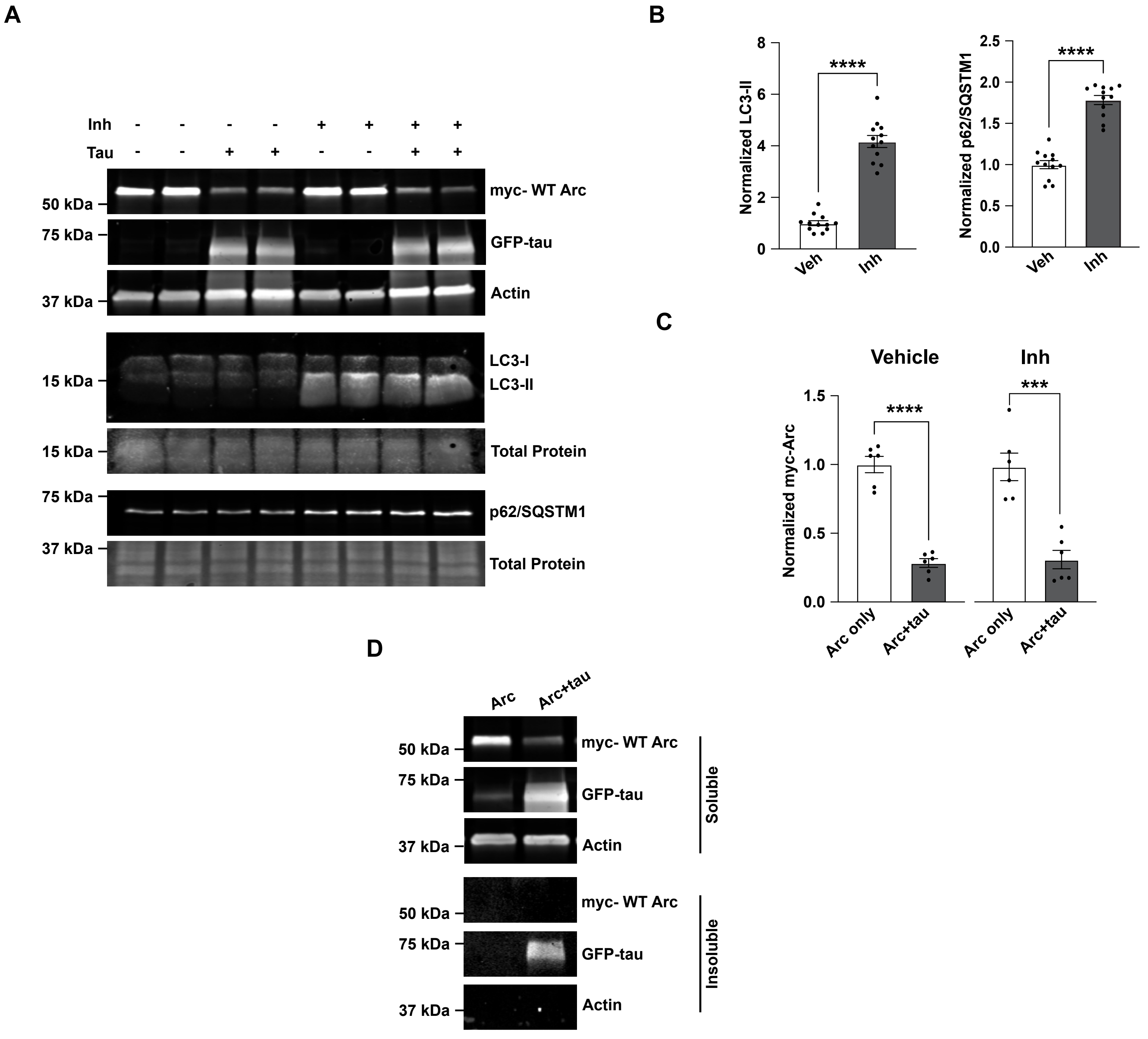
